# Virulence-linked adhesin drives mutualist colonization of the bee gut via biofilm formation

**DOI:** 10.1101/2024.10.14.618124

**Authors:** Patrick J. Lariviere, A. H. M. Zuberi Ashraf, Isaac Gifford, Sylvia L. Tanguma, Jeffrey E. Barrick, Nancy A. Moran

## Abstract

Bacterial biofilms are stable multicellular structures that can enable long term host association. Yet, the role of biofilms in supporting gut mutualism is still not fully understood. Here, we investigate *Snodgrassella alvi*, a beneficial bacterial symbiont of honey bees, and find that biofilm formation is required for its colonization of the bee gut. We constructed fifteen *S. alvi* mutants containing knockouts of genes known to promote colonization with putative roles in biofilm formation. Genes required for colonization included *staA* and *staB*, encoding trimeric autotransporter adhesins (TAAs) and *mltA*, encoding a lytic transglycosylase. Intriguingly, TAAs are considered virulence factors in pathogens but support mutualism by the symbiont *S. alvi. In vitro*, biofilm formation was reduced in Δ*staB* cells and abolished in the other two mutants. Loss of *staA* also reduced auto-aggregation and cell-cell connections. Based on structural predictions, StaA/B are massive (>300 nm) TAAs with many repeats in their stalk regions. Further, we find that StaA/B are conserved across *Snodgrassella* species, suggesting that StaA/B-dependent colonization is characteristic of this symbiont lineage. Finally, *staA* deletion increases sensitivity to bactericidal antimicrobials, suggesting that the biofilm indirectly buffers against antibiotic stress. In all, the inability of two biofilm-deficient strains (Δ*staA* and Δ*mltA*) to effectively mono-colonize bees indicates that *S. alvi* biofilm formation is required for colonization of the bee gut. We envision the bee gut system as a genetically tractable model for studying the physical basis of biofilm-mutualist-gut interactions.

## Introduction

Most animals host microbial communities referred to as their microbiomes or microbiota. Host-associated microbiomes may influence host health by contributing to host metabolism, nutrient production, protection against pathogens, and immune system stimulation^1–3^, among other activities. Understanding how microbiome members colonize a host to set up and maintain structures that result in a stable physical association is crucial for investigations into host-symbiont relationships.

Biofilm formation facilitates host colonization both by pathogens^4–11^ and symbionts^12–18^. A biofilm is formed when bacteria initially adhere to a surface, then construct a protective matrix of secreted macromolecules and components from dead cells around living cells, collectively known as a biofilm^19–25^. Deeper mechanistic understanding of the factors driving biofilm formation comes from investigation of biofilm-forming pathogens and symbionts. For example, global transcriptional regulators such as *rpoN* positively regulate biofilm formation in many bacteria^26–36^, presumably in response to environmental signals. Adhesins, on the other hand, such as the multi-protein Type 4 pilus (T4P) complex and the virulence-associated trimeric autotransporter adhesins (TAAs), directly facilitate physical attachment of bacteria to biotic and/or abiotic surfaces^37–42^. Lytic transglycosylases, such as *mltA*, lyse cells and release extracellular DNA (eDNA), a structural component of the biofilm^43,44^. While biofilm formation has been suggested to play a role in host colonization by gut mutualists, its necessity remains uncertain. Additionally, the molecular mechanisms underlying both biofilm formation and colonization by gut mutualists are not yet fully understood.

The honey bee gut microbiome is emerging as a model for investigating mutualist-host interactions. Honey bees contain a conserved core gut microbiome of 5-8 members^45^. One microbiome member beneficial for host health, *Snodgrassella alvi*, colonizes the ileum through biofilm formation^17^, where it contributes to host immune system function^46,47^ and protection against pathogens^48,49^. *S. alvi* is also thought to deplete oxygen in the ileum, to the benefit of other bee gut bacteria^50^. In a transposon insertion sequencing (Tn-seq) study, genes implicated in *S. alvi* biofilm formation were found to promote host colonization^51^. However, because Tn-seq experiments assess mutant fitness in competition against wild type, as opposed to mutant fitness alone, it remains unclear if biofilm formation is *required* for colonization. Further, the mechanism underlying *S. alvi* biofilm formation is unresolved.

Here, we take a reverse genetics approach to determine if putative biofilm formation genes in *S. alvi* are necessary for colonization. We tested knocking out adhesin genes, biofilm-associated transcriptional regulators, and regulators of eDNA production. For genes found to be beneficial or required for colonization, we then assessed their contribution to biofilm formation. Finally, we conducted a deeper investigation into *staA*, a factor required for both biofilm formation and colonization, to gain further mechanistic insight into how *S. alvi* colonizes its bee host.

## Results

### StaA, StaB, and MltA are required for effective host mono-colonization

To test whether biofilm formation is essential for colonization, we engineered strains containing deletions in 15 putative biofilm formation genes, identified as important for colonization (vs. WT) in a previous Tn-Seq experiment^51^. To confirm that WT outcompetes these mutants, we first tested colonization of microbiota-deficient adult bees in a subset of these mutants in the presence of WT. We inoculated bees with a ratio of 90% mutants to 10% WT, to give the advantage to the mutant. We found that, as in the Tn-Seq experiment, mutants could not robustly colonize in the presence of WT (Figs 1A, S1), indicating that they are outcompeted by WT. At least one mutant, Δ*pilT*, colonized the host to a higher degree than did other T4P mutants (Fig 1A), but was still ultimately outcompeted by WT by over 100-fold.

**Figure 1.**
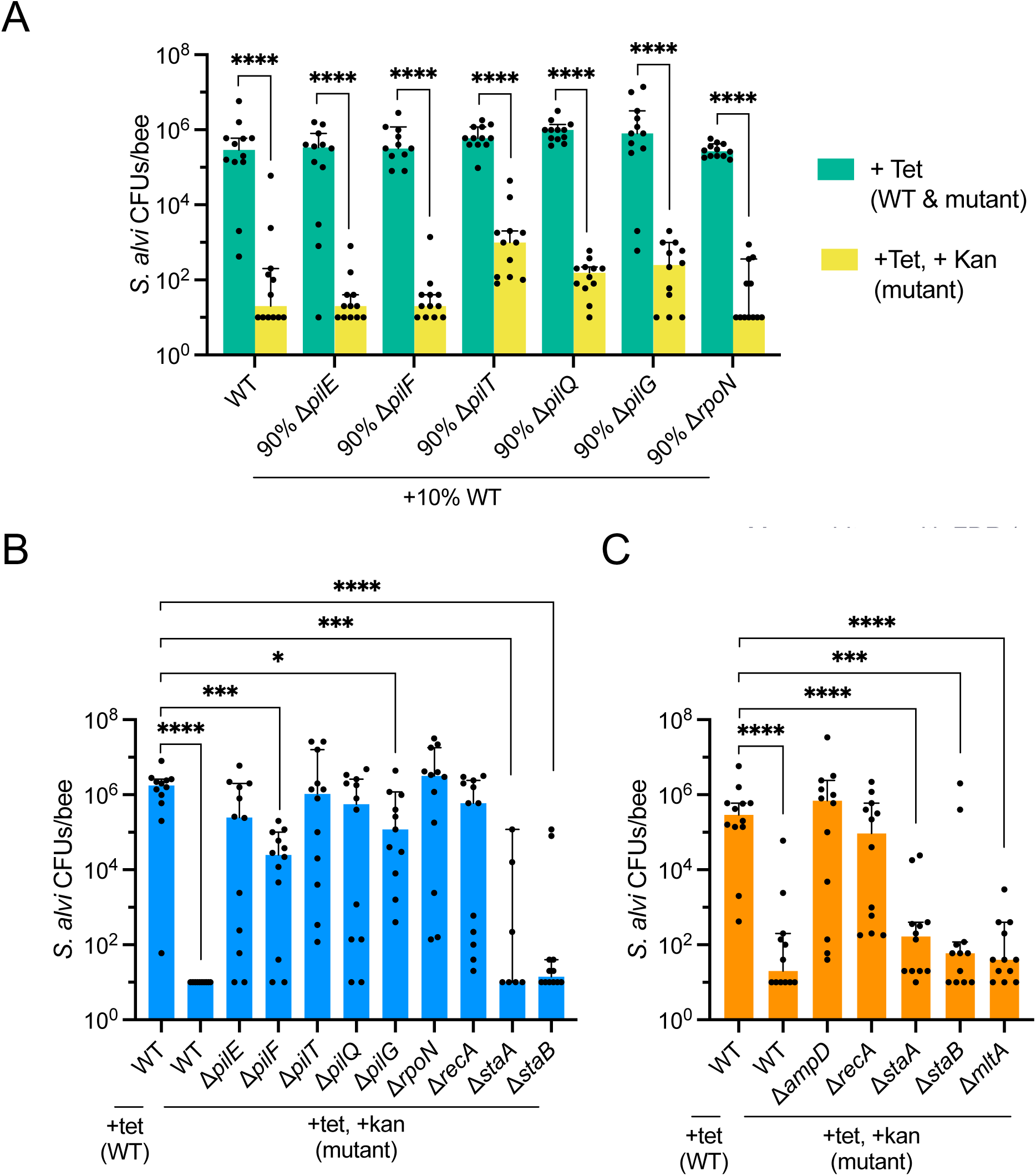
StaA, StaB, and MltA are required for effective host mono-colonization. A. Plot of CFU counts of WT and mutant *S. alvi* cells co-colonized in bees, indicating WT outcompetes the mutant strains. Ilea from bees co-colonized with 10% WT and 90% mutant *S. alvi* were plated on media containing Tet (selects for WT and mutant, green bars) and Tet + Kan (selects for mutant, yellow bars). N = 11-12 biological replicates for each condition. A Shapiro-Wilk test found that not all log_10_-transformed data are normally distributed. Individual data points and group medians with 95% CI are shown. Significant differences between the indicated log_10_-transformed group medians were determined by Mann-Whitney U tests with the two-stage Benjamini, Krieger, & Yekutieli method to correct for multiple comparisons by controlling the FDR (\*\*\*\**q* ≤ 0.0001). B. Plot of CFU counts of *S. alvi* mutants mono-colonized in bees, indicating Δ*staA* and Δ*staB* cannot effectively mono-colonize bees. Ilea from bees mono-colonized with mutant *S. alvi* were plated on media containing Tet + Kan (selects for mutant). Ilea from bees mono-colonized with WT control were plated on both Tet (allows for WT growth) and Tet + Kan (does not allow for WT growth). N = 7-12 biological replicates for each condition. A Shapiro-Wilk test found that not all log_10_-transformed data are normally distributed. Individual data points and group medians with 95% CI are shown. Significant differences between the log_10_-transformed medians of WT (+Tet) and all other groups were determined by Mann-Whitney U tests with the two-stage Benjamini, Krieger, & Yekutieli method to correct for multiple comparisons by controlling the FDR (\**q* ≤ 0.05, \*\*\**q* ≤ 0.001, \*\*\*\**q* ≤ 0.0001). C. Plot of CFU counts of *S. alvi* mutants mono-colonized in bees, indicating Δ*mltA* cannot effectively mono-colonize bees. Δ*staA*, Δ*staB* were also confirmed to not effectively mono-colonize bees. Ilea from bees mono-colonized with mutant *S. alvi* were plated on media containing Tet + Kan (selects for mutant). Ilea from bees mono-colonized with WT control were plated on both Tet (allows for WT growth) and Tet + Kan (does not allow for WT growth). N = 12 biological replicates for each condition. A Shapiro-Wilk test found that not all log_10_-transformed data are normally distributed. Individual data points and group medians with 95% CI are shown. Significant differences between the log_10_-transformed medians of WT (+Tet) and all other group were determined by Mann-Whitney U tests with the two-stage Benjamini, Krieger, & Yekutieli method to correct for multiple comparisons by controlling the FDR (\*\*\**q* ≤ 0.001, \*\*\*\**q* ≤ 0.0001).

We then tested if biofilm formation is required for host colonization by mono-colonizing bees with the mutant alone. Unexpectedly, we found that all the T4P mutants were able to colonize to varying degrees, to the same final cell number as wild type in some cases (Figs 1B, S2). Therefore, while the T4P is beneficial for colonization in the presence of competitors, it is not required in mono-colonization. By contrast, mutants with deletions in two TAAs, *staA* and *staB*, and one lytic transglycosylase, *mltA*, were unable to effectively mono-colonize (Figs 1B, 1C, S2, S3). For these strains, most bees had little to no bacterial growth. A couple of bees in each of these groups showed some colonization, which we suspect was due to suppressor mutations compensating for the Δ*staA* phenotype within some bees. Repeat experiments of Δ*staA* and Δ*staB* mono-colonization (Figs 1C, S3, S4) confirmed nearly complete loss in colonization compared to WT. From these results, we conclude that StaA, StaB, and MltA are required for effective mono-colonization.

To test if ineffective mono-colonization was due to an overall reduction in cell growth rate, we measured CFUs and growth rate for a few *in vitro* grown mutant strains (Fig S5). For Δ*staA*, while growth rate was slightly reduced in Insectagro (Fig S5C), *in vitro* final CFU counts and growth rate were actually higher than WT in Columbia media (Fig S5A, B). For Δ*rpoN*, growth rate was slightly reduced (Fig S5E), but *in vitro* CFU counts were comparable to WT (Fig S5A). For Δ*mltA*, *in vitro* CFU counts were reduced (Fig S5A), but growth rate was similar to WT for 2 out of 3 replicates (Fig S5D). Taken together, these results do not reveal substantial growth differences between the mutants tested and WT, indicating that growth rate alone does not account for the observed differences in colonization among these mutants compared to WT.

### StaA and MltA are required for biofilm formation

To test if mutants deficient in colonization were able to form biofilm, we quantified biofilm formation using a crystal violet assay^51–53^. We found that the T4P mutants had significantly reduced, but non-zero biofilm production (Fig 2A-C). These results align with previous reports in *S. alvi* that biofilm formation in Δ*pilF* and Δ*pilG* strains is somewhat reduced^51,52^. Biofilm formation was completely abolished in Δ*staA* and Δ*mltA*, on the other hand (Fig 2). These results provide evidence that biofilm formation is required for colonization, as these were two of the mutants unable to effectively mono-colonize bees (Fig 1). Intriguingly, Δ*staB* cells still produced biofilm *in vitro*, suggesting that StaB’s colonization dependence stems from another function. Biofilm formation was also substantially reduced in Δ*rpoN*, suggesting that RpoN may play an upstream transcriptional regulation role in biofilm formation (Fig 2D-F). Finally, strains containing deletions in other factors including *recA*, *recJ, tspA*, *ampD, amsE*, *wcwK* still produced biofilm (Fig 2E-F). However, these strains had reduced biofilm-to-growth ratios compared to WT (Fig 2D), indicating that other factors likely contribute to biofilm production. Taken together, these results highlight StaA and MltA as essential for biofilm formation, and implicate other factors, particularly the T4P, as important for biofilm formation.

**Figure 2.**
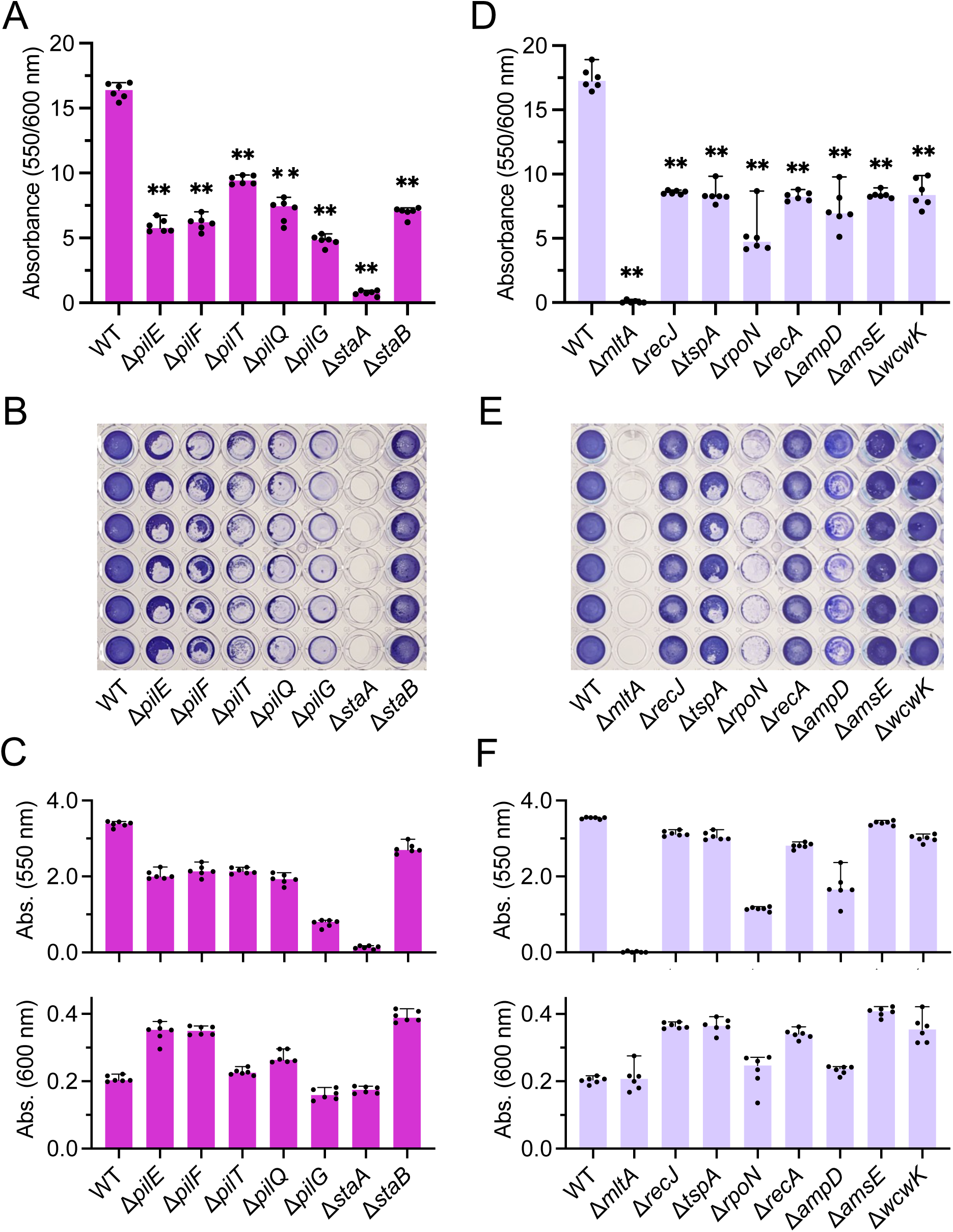
StaA and MltA are required for biofilm formation. A. Plot quantifying cell growth-normalized biofilm formation, indicating biofilm formation is decreased in most strains and abolished in Δ*staA*. N = 6 for each condition. A Shapiro-Wilk test found that not all data are normally distributed. Individual data points and group medians with 95% CI are shown. Significant differences between the medians of WT and other groups were determined by Mann-Whitney U tests with the two-stage Benjamini, Krieger, & Yekutieli method to correct for multiple comparisons by controlling the FDR (\*\**q* ≤ 0.01). B. Image of 96-well plate stained with crystal violet, indicating biofilm formation is abolished in Δ*staA* and reduced to varying degrees in other mutants. C. Plots quantifying non-normalized biofilm formation (top) and cell growth (bottom), demonstrating Δ*staA* has near normal growth, but does not form biofilm. Other mutants produce varying degrees of biofilm, with OD_600_ values similar to or higher than WT. N = 6 for each condition. A Shapiro-Wilk test found that not all data are normally distributed. Individual data points and group medians with 95% CI are shown. D. Plot quantifying cell growth-normalized biofilm formation, indicating biofilm formation is abolished in Δ*mltA* and decreased to varying degrees in other mutants. N = 6 for each condition. A Shapiro-Wilk test found that not all data are normally distributed. Individual data points and group medians with 95% CI are shown. Significant differences between the medians of WT and other groups were determined by Mann-Whitney U tests with the two-stage Benjamini, Krieger, & Yekutieli method to correct for multiple comparisons by controlling the FDR (\*\**q* ≤ 0.01). E. Image of 96-well plate stained with crystal violet, indicating biofilm formation is abolished in Δ*mltA*, reduced in some mutants, and close to WT levels in others. F. Plots quantifying non-normalized biofilm formation (top) and cell growth (bottom), demonstrating Δ*mltA* has normal growth, but does not form biofilm. Some mutants have decreased or WT-like biofilm formation, with WT-like or better growth. N = 6 for each condition. A Shapiro-Wilk test found that not all data are normally distributed. Individual data points and group medians with 95% CI are shown.

Secondary mutations have been detected in engineered *S. alvi* strains, including some used in this study. Often these affect the predicted autotransporter-encoding gene SALWKB2_RS07515^52^. To evaluate this mutation’s impact on biofilm formation, we assessed biofilm formation in PL75, which has a frameshift mutation in SALWKB2_RS07515, but intact biofilm genes. We saw no significant difference in biofilm formation in PL75 compared to WT (Fig S6), indicating mutations in SALWKB2_RS07515 do not confound our other biofilm formation data.

Finally, we assessed biofilm formation by *S. alvi* in multiple media types and found that Insectagro (Fig 2), but not BHI (Fig S7) is conducive to biofilm formation. This was not due to differences in growth rates, as cell growth was unaffected by change in media for WT *S. alvi* and multiple mutants (Fig S7F, I). This result indicates that nutrient/solute availability can influence *S. alvi* biofilm formation, independent of cell growth.

### StaA promotes auto-aggregation in *S. alvi* cells

While performing the crystal violet assay, we noticed that WT forms clumps in liquid media, whereas Δ*staA* cells do not. To validate this observation, we assessed WT and Δ*staA* cell sedimentation. We found that, whereas WT rapidly sediments, Δ*staA* does not (Fig 3A, B). These results led us to hypothesize that, as an adhesin-containing protein, StaA facilitates auto-aggregation. To test this hypothesis, we first performed DIC microscopy to visualize any auto-aggregation of resuspended WT or Δ*staA* cells. Strikingly, large clumps were observed for WT but not Δ*staA* cells (Fig 3C). We subsequently performed flow cytometry on OD_600_-normalized WT and Δ*staA* cells to quantify differences in aggregation. We observed a particle population with increased size (forward scattering) for WT compared to Δ*staA* (Fig 4C), confirming that large cell clumps are less prevalent in Δ*staA* populations. In all, these results indicate that StaA promotes auto-aggregation of *S. alvi* cells.

**Figure 3.**
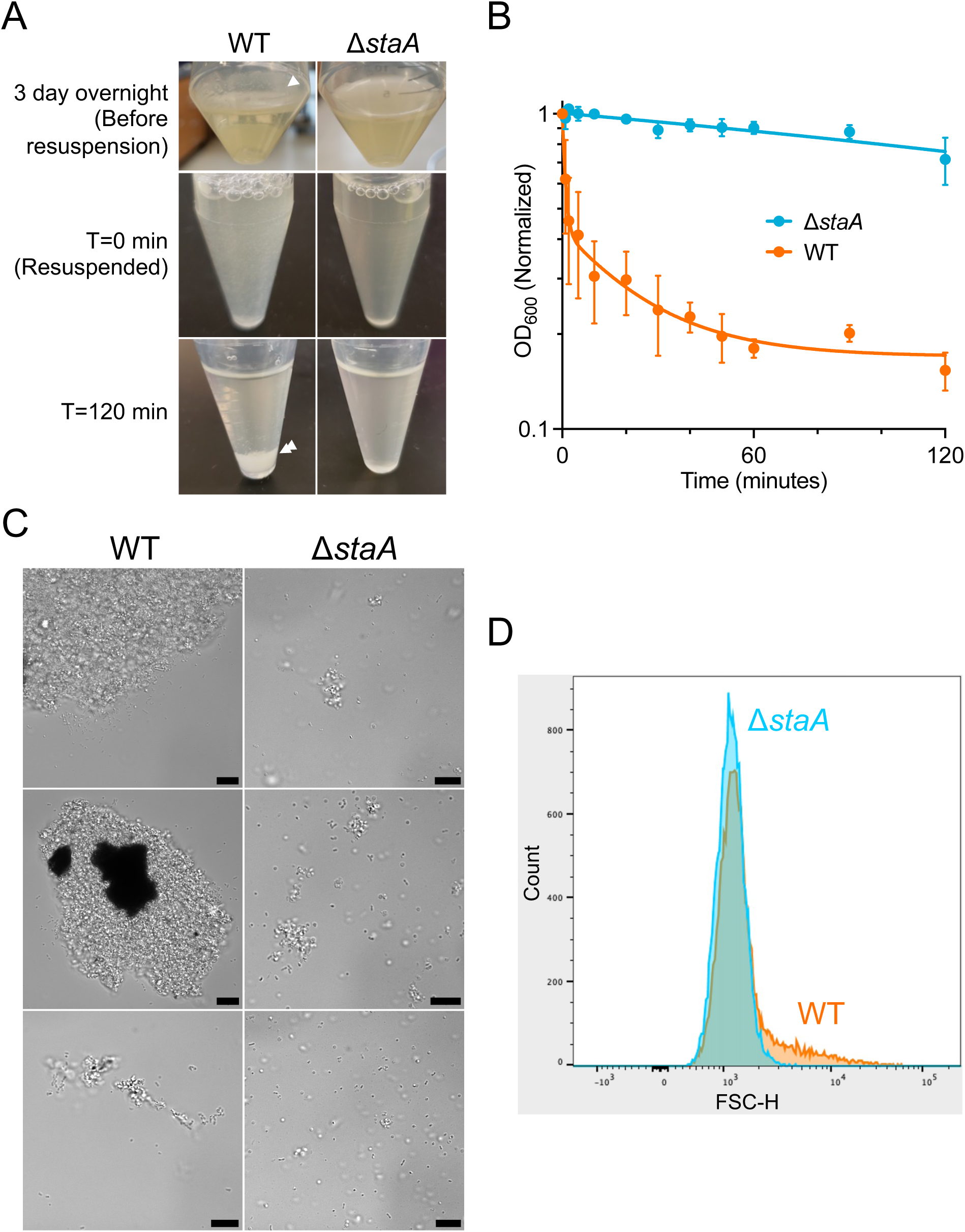
StaA promotes auto-aggregation in *S. alvi* cells. A. Images of tubes containing WT or Δ*staA* before (top) and after resuspension (middle) that were allowed to sediment (bottom), demonstrating that WT but not Δ*staA* noticeably sediments within 120 min. WT cells form a biofilm prior to resuspension (top left, single arrow), briefly remain resuspended (middle left), and sediment after resuspension (bottom left, double arrow). Δ*staA* cells primarily remain planktonic before (top right) and after (middle and bottom right) resuspension. B. Plot of cell density (normalized OD_600_, log scale) taken from the top of resuspended cell cultures at different timepoints post-resuspension. WT (blue) quickly sediments, whereas Δ*staA* (orange) does not. N = 3 biological replicates for each condition. Trendline = Non-linear best fit (two-phase decay model); data points = means of biological replicates; error bars = SD. C. DIC microscopy images of WT (resuspended from biofilm, left) and Δ*staA* (planktonic, right) cells after growth in liquid culture. Micrographs demonstrate that WT cells form large auto-aggregates, whereas Δ*staA* cells do not. Specifically, WT cells form larger (top left, middle left) and smaller (bottom left) aggregates, whereas Δ*staA* cells are planktonic (right) or found in smaller aggregates (top right, middle right). Rows depict different fields of view from the same experiment. Scale bar = 10 μm. D. Plot of forward scatter of populations of WT (orange) and Δ*staA* (blue) cells measured during flow cytometry. The WT forward scatter distribution has a right shoulder absent in Δ*staA*, indicating that WT cells form larger aggregates than Δ*staA* cells.

**Figure 4.**
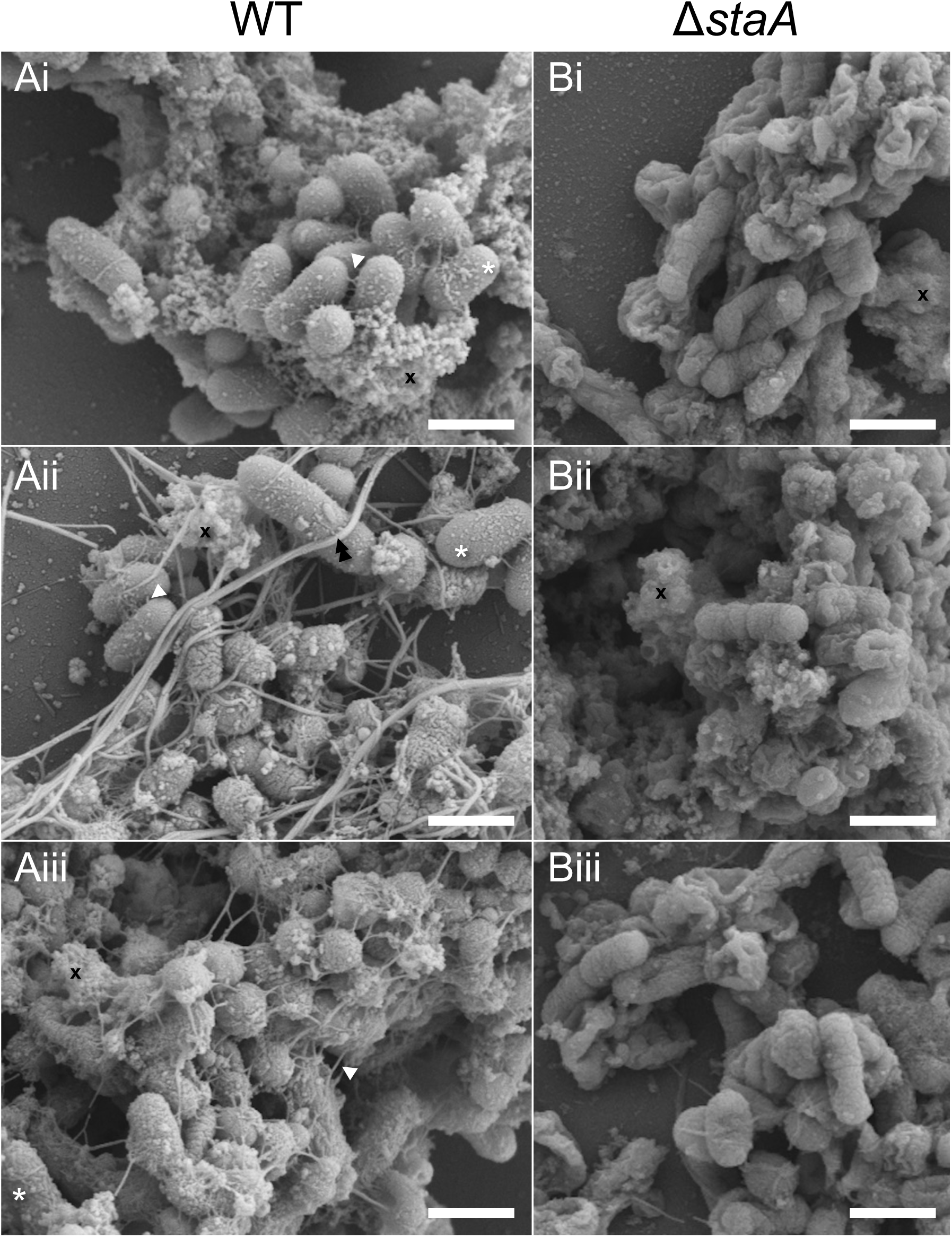
StaA promotes formation of cell-cell connections. SEM images of WT (Ai-Aiii) and Δ*staA* (Bi-Biii) cells. Short range cell-cell connections (white arrow) and cell surface knobs (white asterisk) are present in WT cells, but are largely absent in Δ*staA* cells. Longer strands (black double arrow) are presumed to be biofilm matrix material. EPS (black x) is observed in both WT and Δ*staA* cells. Rows represent different fields of view from the same experiment. Scale bar = 1 μm.

### StaA promotes formation of cell-cell connections

To further characterize how StaA promotes auto-aggregation, we visualized WT and Δ*staA* cells at higher resolution. We performed scanning electron microscopy (SEM) on resuspended WT and Δ*staA* cells grown in liquid culture (Fig 4). EPS (extracellular polymeric substance) was observed in both populations, though it was more prominent in WT samples (Fig 4). More strikingly, WT cells were observed to form clusters linked by short cell-cell connections (Fig 4A). Individual WT cells had a knob-like surface texture (Fig 4A). Δ*staA* cells, on the other hand, lacked cell-cell connections and had smooth cell surfaces lacking the knobs seen in WT (Fig 4B). Such cell-cell connections and surface knobs have been associated with TAAs in other species^54,55^ (see discussion). We therefore conclude that StaA promotes formation of cell-cell connections, which we propose drives auto-aggregation.

### StaA, StaB, and StaC are massive TAAs

To characterize *S. alvi* TAA gene architecture, we made protein sequence diagrams with domain annotations of the three paralogs present in WT: *staA, staB,* and *staC* (Fig 5A). The *staC* gene is a homolog of *staA* and *staB* but was not found to promote colonization in the prior transposon mutagenesis screen^51^, which is why it was not tested for colonization and biofilm formation in this study. Similar to other YadA-like TAAs, *staA* and *staB* each encode a signal peptide (presumed to drive localization to the *sec* translocon), adhesive head region, stalk and neck repeats, and an outer-membrane anchor (Fig 5A). Remarkably, *staA* and *staB* were found to be substantially longer than previously described TAAs, including YadA (*Yersinia*), NadA (*Neisseria*, a relative of *Snodgrassella*), and UpaG (*Escherichia coli*). Increased gene size is due, in large part, to the 27+ head and neck repeats found in the *S. alvi* TAAs. (Fig 5A).

**Figure 5.**
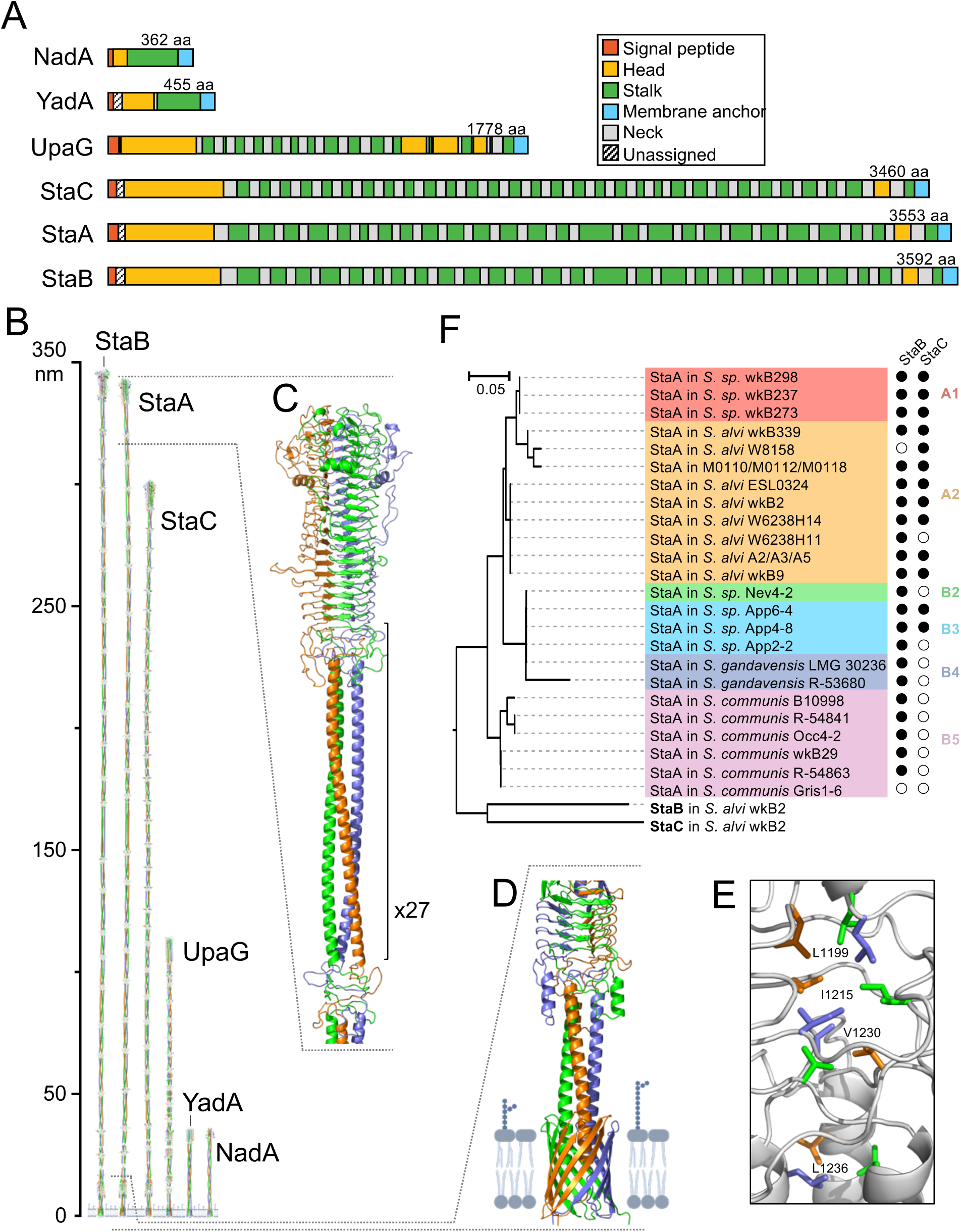
StaA and StaB are massive TAAs that are conserved in *Snodgrassella*. A. Protein domain diagrams of TAAs from *S. alvi* (StaA/B/C) and other bacteria (NadA, YadA, UpaG), demonstrating the *S. alvi* TAAs are encoded by large genes. Genes and domains are depicted to scale. The number of amino acids (aa) encoded by each gene is shown above each gene. Domain colors are indicated in the key. B. Predicted protein structures of TAAs from *S. alvi* (StaA/B/C) and other bacteria (NadA, YadA, UpaG), demonstrating the *S. alvi* TAAs are massive proteins. Within individual structures, each monomer has a single color (red, blue, or green). Membrane anchor domains are shown inserted in the outer membrane. Structures are depicted to scale. C. Zoomed in view of the StaA head, neck, and stalk domains. The neck + stalk superdomain repeats 27 times, as indicated, throughout the length of the protein. Each monomer has a single color (red, blue, or green). D. Zoomed in view of the StaA membrane anchor, shown inserted into the outer membrane. Each monomer has a single color (red, blue, or green). E. Zoomed in view of a neck domain hydrophobic core, highlighting that the neck pincushions three monomers together. Putative interacting residues are shown and labeled, with colors (red, blue, or green) representing distinct monomers. Other neck residues are shown in white (for all 3 monomers). F. Phylogenetic tree of the StaA anchor domain in *Snodgrassella* (left), indicating StaA is conserved across the genus. Annotation of presence or absence of StaB/StaC in the indicated species (right), indicating conservation of StaB in *Snodgrassella*. Tree is drawn to scale, as indicated. StaB and StaC from wkB2 root the tree. *Snodgrassella* taxonomic groups are denoted by colors and are labeled (*Apis*-specific species: A1, A2; *Bombus*-specific species: B2, B3, B4, B5) according to the subclades previously identified on the basis of 254 core proteins^61^. Black circle: StaB/StaC ortholog present in the indicated species; White circle: StaB/StaC ortholog absent in the indicated species.

Next, we predicted the structures of the *S. alvi* TAAs to gain further insight into their functions (Fig 5B). StaA/B/C are massive in size compared to previously described TAAs (Fig 5B). In fact, StaA/B/C are predicted to be nearly 3x the length of UpaG, the largest TAA that has been visualized^55^. We therefore hypothesize that *S. alvi* TAA length specifically evolved to overcome physical barriers to host colonization in the bee ileum (see discussion).

Domain level analysis revealed similarities to other YadA-like TAAs. The signal sequence is presumably cleaved and was omitted from the structure prediction. Similar to other YadA-like TAAs, StaA/B/C trimerize across all domains (Fig 5B). The N-terminal YadA-like head, which has been shown to be the domain primarily responsible for adhesion in other organisms^56–60^, is composed of a trimeric beta helix (Fig 5C). C-terminal to the head is a globular neck domain, which pincushions the three monomers with a hydrophobic core (Fig 5E), and a longer stalk domain composed of a trimeric coiled-coil (Fig 5C). This stalk + neck superdomain repeats 26 additional times to compose the bulk of the protein, with stalks varying in length (Fig 5B,C). Alignment of the 27 stalk + neck repeats revealed conserved residues in both the neck and stalk (Fig S8). A phylogenetic tree of these repeats highlights relatedness between different copies (Fig S8B), suggesting that the repeats arose due to multiple duplication events. Finally, a smaller YadA-like head domain is found close to the C-terminus of the protein (Fig 5D), just before the beta barrel outer membrane anchor domain (Fig 5D).

StaB has a similar overall structure to StaA, with domain-level differences. Notably, the StaB head domain contains an unsatisfied, charged loop sticking out into space that is absent in StaA (Fig S9). Such a loop is suggestive of a ligand binding site, whose function could be to interact with a host factor. Accordingly, we hypothesize that StaB could be required for colonization due to putative binding to a host factor.

### StaA and StaB are conserved among *Snodgrassella*

To generate insight into the potential conservation of StaA/StaB-mediated colonization, we identified orthologs of StaA/StaB/StaC in the bee microbiome via BLAST. Numerous orthologs were found in other *Snodgrassella* species, but not in other bee gut symbionts. We constructed a phylogenetic tree for StaA, for which orthologs were broadly conserved across *Snodgrassella* (Fig. 5F). Clusters largely correspond to species groups previously found in a phylogeny constructed on the basis of 254 core genes^61^, supporting a deep history of this protein in *Snodgrassella*. StaA was present in all species groups, from both honey bees (genus *Apis*) and bumble bees (genus *Bombus*), with the exception of one *Bombus* group (B1^61^) (Fig 5F). StaB was similarly present in most *Snodgrassella* species, whereas StaC was primarily found in *Apis*-specific species (Fig 5F). Finally, StaA orthologs were found to be more closely related to one another than to StaB or StaC (Fig 5F), indicating that the duplications giving rise to paralog copies occurred prior to the diversification of these symbionts and their hosts, over 80 million years ago^62^. In all, the broad presence of StaA/StaB within the genus suggests that TAA-mediated host colonization within *Snodgrassella* is conserved.

To identify conserved motifs within StaA, we aligned orthologs from within the genus. We found particularly high conservation within the N-terminal region, which allowed us to identify boundaries of the signal sequence (Fig S10A). We performed a similar analysis to determine the signal sequence for StaB (Fig S10B), which was similar to that of StaA. StaA and StaB therefore each have conserved, related signal sequences. These results suggest that inner membrane localization of StaA/StaB is conserved across the genus, providing further evidence for conservation of TAA function in *Snodgrassella*.

Additionally, we assessed conservation of StaA/StaB length. We found StaA/StaB length was relatively conserved (∼3600 amino acids) in the *Apis*-colonizing *Snodgrassella* groups A1, A2 and the *Bombus*-colonizing group B5 (Table S1). TAA length was shorter (∼1950 amino acids) in the *Bombus*-colonizing groups B2, B3, and B4 (Table S1), resulting from the absence of multiple stalk+neck repeats. Thus, TAA length is variable across host genera.

### The *S. alvi* biofilm is mildly protective against bactericidal antimicrobials

Biofilms have been shown to protect bacteria from external stressors such as antimicrobials^22,63–67^. To determine if *S. alvi* biofilms can similarly protect against antimicrobials, we exposed biofilm competent (WT) and biofilm deficient (Δ*staA*) cells to the bactericidal antibiotic gentamicin (Fig 6A). We found Δ*staA* cells had steeper reduction in growth relative to antibiotic concentration compared to WT (Figs 6A, S11). This result indicates that the *S. alvi* biofilm has a mildly protective effect against antimicrobials. We next sought to determine if the biofilm could protect against a physiologically relevant antimicrobial that *S. alvi* might encounter in its host. The honey bee produces a cocktail of antimicrobial peptides, including apidaecin 1B, which is present in the bee gut^46^. We found that Δ*staA* was more susceptible to apidaecin 1B than was WT at higher concentrations (Figs 6B, S11). This result suggests that, in addition to facilitating initial colonization, the *S. alvi* biofilm could play a role in protecting against stresses such as host-produced antimicrobial peptides.

**Figure 6.**
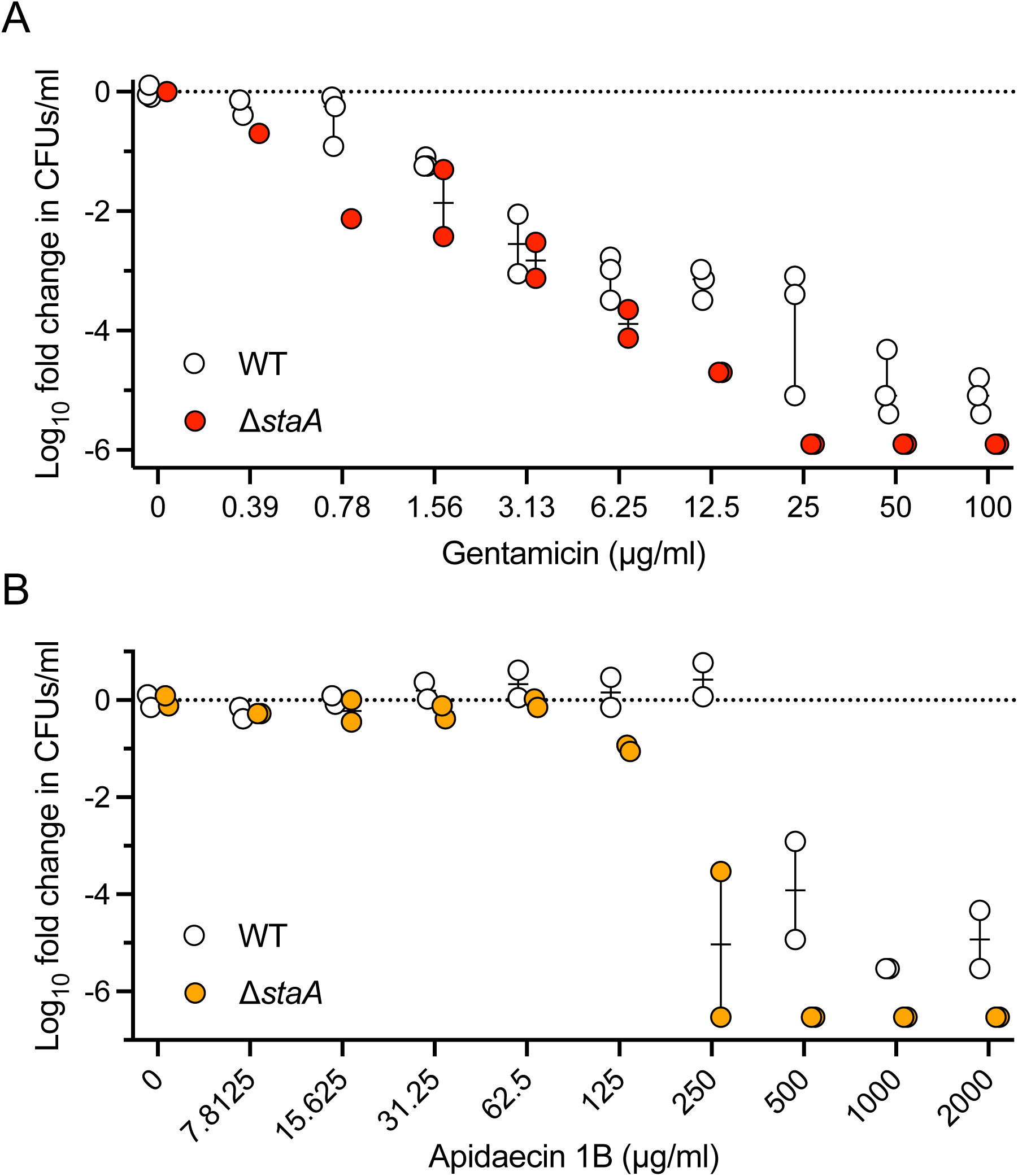
The *S. alvi* biofilm is mildly protective against antimicrobials. A. Plot of log_10_ fold change in CFUs/ml of WT (white) or Δ*staA* (red) exposed to increasing concentrations of gentamicin, indicating that Δ*staA* cells are slightly more sensitive to higher concentrations of gentamicin. Fold change is normalized to baseline CFUs/ml of cells not exposed to gentamicin. Dotted line (fold change baseline): 0-fold change in log_10_ normalized CFUs/ml. For each group, individual data points and medians with 95% CI are shown. N = 1-3 biological replicates per condition. B. Plot of log_10_ fold change in CFUs/ml of WT (white) or Δ*staA* (orange) exposed to increasing concentrations of apidaecin 1B, indicating that Δ*staA* cells are more sensitive to higher concentrations of apidaecin 1B. Fold change is normalized to baseline CFUs/ml of cells not exposed to gentamicin. Dotted line (fold change baseline): no change in log_10_ normalized CFUs/ml. For each group, individual data points and medians with 95% CI are shown. N = 2 biological replicates per condition.

## Discussion

Here we report that biofilm formation is required for effective colonization of honey bee guts by the host-specialized mutualist *S. alvi*. We find that the TAAs StaA and StaB are necessary for colonization. StaA, but not StaB, is required for *in vitro* biofilm formation. This discrepancy suggests StaA acts as a generally adhesive protein that binds non-specifically, whereas StaB has specificity for a host factor. We therefore propose a model whereby StaA drives biofilm formation through auto-aggregation and non-specific surface binding to facilitate effective host colonization by *S. alvi* (Fig 7). In this model, StaB promotes colonization through specific binding to one or multiple unidentified host factors (Fig 7), possibly through its charged head loop (Fig S9). We propose that StaA/StaB-mediated colonization is conserved in *Snodgrassella*, since StaA/StaB orthologs are found throughout the genus.

**Figure 7.**
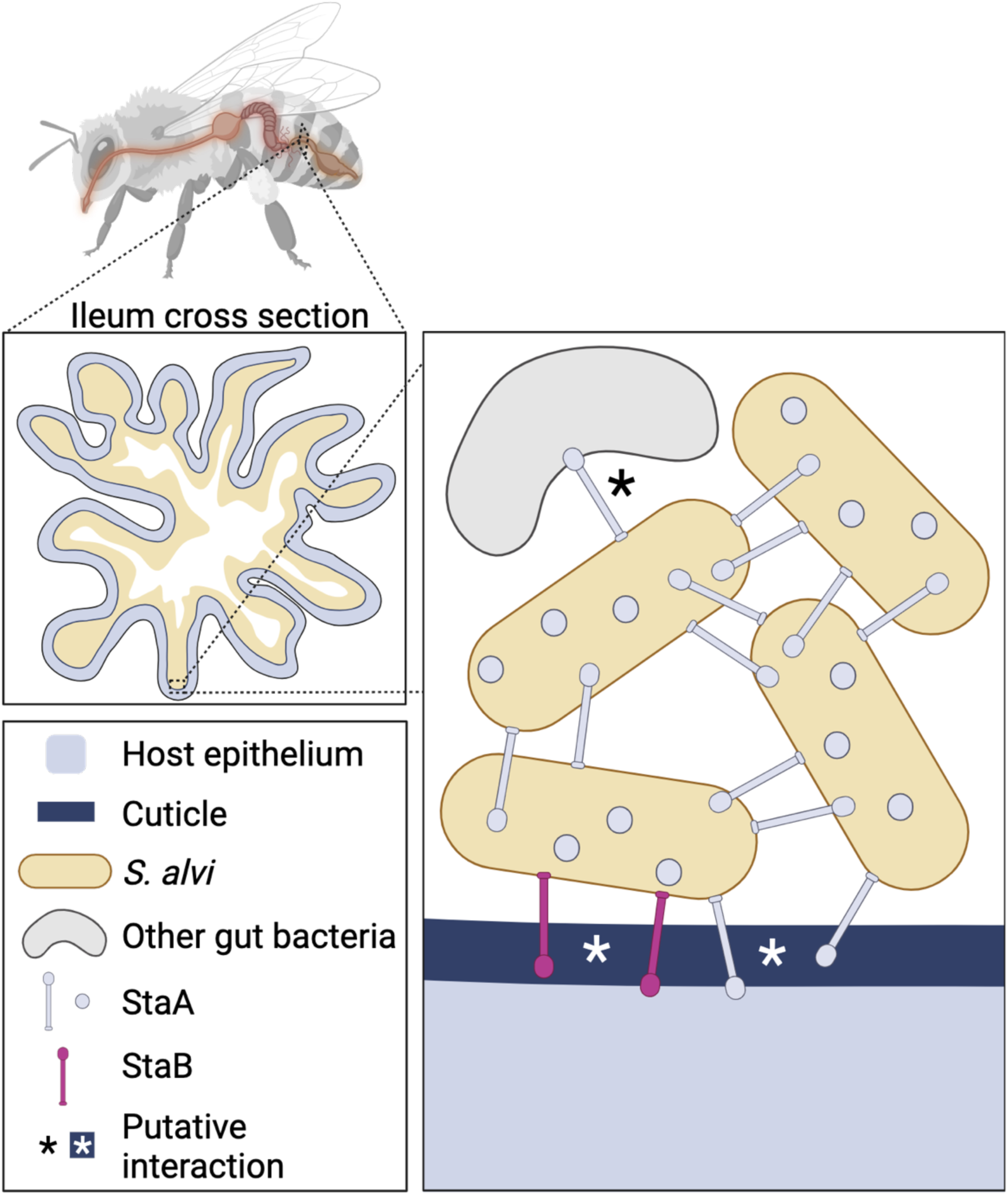
Model for StaA/StaB-mediated host colonization. Cartoon of model for StaA/StaB-dependent colonization of the bee gut by *S. alvi*. Both StaA and StaB are required for effective colonization. As a biofilm-forming adhesin, StaA mediates *S. alvi* auto-aggregation (right) and putatively interacts with host epithelium or cuticle and other gut bacteria (right). As a non-biofilm forming adhesin, StaB is hypothesized to interact with host epithelium or cuticle (right). The host-microbe interaction panel (right) is an inset of a cross section of the ileum (middle left), itself an inset of the bee gut (top left). Key of symbols is shown (bottom left).

TAAs have been shown to be important for biofilm formation^68^, host adherence/colonization^69,70^, or both^60,71–79^ in many Gram-negative bacteria, including *Yersinia*, *Bartonella*, *Neisseria*, *Salmonella*, *E. coli*, *Acinetobacter baumanii*, and others. TAAs facilitate host cell interaction by binding to host proteins. Many TAAs specifically interact with host extracellular matrix proteins, as BadA from *B. henselae* does with collagen and fibronectin^77,80^. Distinct charged loops in the YadA heads from *Yersinia enterocolitica* and *Yersinia pseudotuberculosis* specify binding to host vitronectin^59^ and fibronectin^60^, respectively. These findings lend credibility to our hypothesis that StaB’s charged head loop determines host factor specificity. Meanwhile, at least one TAA (AtaA in *Acinetobacter*) has been characterized as displaying non-specific stickiness^81^. Further work remains to identify any StaB host-binding partners and to determine whether StaA displays any binding specificity. TAAs also induce auto-aggregation, which can improve biofilm formation by physically clumping cells together ^68,71,73–76,82^. However, it is unclear if the TAA binding partner during auto-aggregation is a TAA on another cell, the cell body itself, or both. StaA similarly induces auto-aggregation, though the binding partner facilitating this interaction is also unknown.

Structurally, there are key similarities between the TAAs in *S. alvi* and those previously described in the literature. At a cellular level, our SEM data suggests that StaA promotes cell-cell connections and forms surface knobs on *S. alvi*. TAAs in other bacteria have been found to form similar structures. For example, BadA facilitates short-range cell-cell connections^54^ and SadA forms knob-like surface textures^55^. These data suggest StaA is likely directly responsible for similar structures in *S. alvi*, though further work remains to verify this. At a molecular level, StaA and StaB contain all of the domains found in other YadA-like TAAs.

Two key differences distinguish the *S. alvi* TAAs from those previously described. First, most TAAs described in the literature belong to pathogens, including opportunistic pathogens^68,83–88^. Accordingly, TAAs are typically referred to as virulence factors^83,85,86,89^. Our findings of TAA utilization by *S. alvi*, a mutualist, necessitates reclassification of TAAs as more general colonization factors. Second, StaA and StaB are substantially larger than previously visualized TAAs^55^, sparking questions about how and why these TAAs are so large.

*S. alvi* TAAs appear to be long due to both domain duplications and optimization for domain length. StaA and StaB have more than twice as many neck + stalk domains as UpaG, indicating more domain duplication events in the *S. alvi* TAAs. Oddly, we initially observed the *staA*/*staB* genes to be twice as long as *upaG*, but the protein structures to be 3x longer, indicating increased length is not simply due to more domain repeats. Closer examination revealed that a higher proportion of residues in UpaG are found in neck domains compared to StaA/B. StaA/B, on the other hand, have a higher proportion of residues in stalks. Because necks are squat and stalks are long, this residue distribution further increases StaA/B length. These proteins appear to have been optimized for length.

Why might StaA/B have evolved greater length despite the potential for higher metabolic burden from producing massive proteins? StaA/B may be required to cross a physical barrier in order to adhere to the correct host surface. Indeed, in honey bees, *S. alvi* cells are separated from the ileum epithelium by an approximately 200 nm thick, porous cuticle^90,91^. Intriguingly, the wkB2 StaA/B structures predict protein lengths of ∼350 nm, long enough to traverse cuticle pores in honey bees. We hypothesize that StaA/B traverse the cuticle to bind to one or more host factors in the epithelium. Alternatively, StaA/B may traverse other entities such as eDNA to bind the cuticle, or may bind both the cuticle and epithelium. Interestingly, compared to wkB2, StaA/B are shorter in the bumblebee-colonizing *Snodrassella* groups B2, B3, and B4 (Table S1). On the other hand, StaA/B from group B5 were similar in length to the wkB2 orthologs (Table S1). Whether the TAAs from all of these groups are long enough to traverse the bumblebee cuticle remains unclear, as data characterizing how cuticle thickness varies among bee genera could not be readily found.

In addition to StaA and StaB, *S. alvi* contains a third long TAA, StaC, which was not required for colonization in the TnSeq experiments^51^. Potentially StaC, while not required, reinforces the functions of StaA/B or is necessary for other processes such as interactions with other bee gut microbes or transmission between bees within a hive.

The T4P was also found to be important for biofilm formation and colonization. In other bacteria, the T4P is a multifunctional machine involved in biofilm formation, motility, and natural competence^92,93^. We found that the T4P contributes to, but is not required for, biofilm formation in *S. alvi*. Likewise, we found that the T4P promotes host colonization (as previously described^51^) but is not essential for colonization. T4P loss might partially reduce the number of adherent cells in the biofilm, a disadvantage that can be overcome absent competition, but not in the presence of more adherent WT cells. Interestingly, the T4P deletion mutants with the highest level of mono-colonization (Δ*pilT*, the retraction ATPase) also had the strongest biofilm formation, further evidence that biofilm formation is important for colonization. Finally, the ability of even a weak biofilm forming mutant (Δ*pilG*) to mono-colonize suggests that even weak *in vitro* biofilm formation is sufficient for *in vivo* colonization.

In addition to adhesins, other factors were found to be important for *S. alvi* biofilm formation and/or colonization. The lytic transglycosylase MltA was required for effective mono-colonization and biofilm formation. Biofilm formation was slightly reduced in another lytic transglycosylase deletion mutant, Δ*ampD*. MltA and AmpD have been shown to promote lysis and thereby eDNA release in other bacteria^43,44^. Since eDNA is a biofilm matrix component^94^, deletion of these genes would be expected to reduce biofilm production in *S. alvi*. Another factor implicated in biofilm formation in other bacteria is *rpoN*, a nitrogen-sensing transcriptional regulator^26–36^. In *S. alvi*, Δ*rpoN* cells had reduced biofilm formation, but were able to mono-colonize bees. As with the *pilG* deletion, these data suggest that even weak biofilm formation allows for some colonization. Finally, Δ*recA* cells had reduced biofilm formation and colonized the host in an all-or-none binary fashion. RecA has been linked to biofilm formation in other Gram-negative bacteria, though as a negative regulator^95,96^. It is therefore unclear how *recA* deletion leads to reduced biofilm formation in *S. alvi*, particularly given that Δ*recA S. alvi* cells do not have otherwise reduced fitness *in vitro*^52^.

The ability of biofilms to protect against physical, biological, and chemical stress, including antibiotics, has been well described^22,63–67^. A protective shell composed of dead cells^25^ and EPS^22^ acts as a diffusion barrier, increasing the concentration of antibiotics required to inhibit bacterial growth^63^. We show here that the *S. alvi* biofilm is mildly protective against antimicrobials, including apidaecin 1B. Apidaecin 1B is a bee-produced AMP present in the hemolymph and ileum, which is active against *S. alvi* at higher concentrations^46^. It is therefore plausible that, in addition to facilitating initial host-adherence, the *S. alvi* biofilm buffers against the host immune system.

In all, we demonstrate here that biofilm formation is necessary for colonization of honey bee guts by the mutualist *S. alvi*. *S. alvi* colonization is primarily TAA-mediated, though other factors, including the T4P, contribute. Future work aims to elucidate the interactions between *S. alvi* adhesins and the host at a molecular level.

## Methods

### Bacterial cell culture

Bacterial strains used in this study are listed in Table S2 (Strains tab). Unless otherwise specified, *S. alvi* cells were grown on Columbia agar plates + 5% sheep’s blood ^52,97,98^ (solid media) and in Insectagro DS2 (Corning; liquid media). (*S. alvi* was grown in BHI liquid media in two experiments, where indicated). *S. alvi* solid and liquid cultures were typically incubated for 2-3 days at 35°C, 5% CO_2_. Where indicated, *S. alvi* was grown with antibiotics at the following indications: Kanamycin (Kan; 25 μg/ml); Tetracycline (Tet; 7.5 μg/ml); Gentamicin (Gent; 0.39 – 100 μg/ml); Apidaecin 1B (7.8125 – 2000 μg/ml); Spectinomycin (30 μg/ml).

*E. coli* NEB5-alpha (for cloning pPL32 and pPL373) was grown in LB broth and on LB agar, at 37°C. Spectinomycin (60 μg/ml) was added to media when necessary.

### Statistical analysis

Unless otherwise specified, all statistical analyses and graphing were performed using GraphPad Prism 9 (Dotmatics, USA).

### *S. alvi* mutant strain engineering

*S. alvi* mutant strains containing gene knockouts were constructed using a previously described one-step genome engineering approach^52^. Specifically, we generated a short list of genes found to be beneficial for host colonization in a previous TnSeq study^51^ that were also putatively involved in biofilm formation. Next, we designed deletion cassettes containing an antibiotic resistance gene flanked by arms with homology to the gene of interest. Deletion cassettes were commercially synthesized as gene fragments inserted in plasmids, amplified, and electroporated as linear DNA into *S. alvi*. Following overnight recovery, *S. alvi* was plated on selective media. Successful transformants were passaged onto a second plate, and then screened for insertion by PCR. To verify success of the transformations and to detect possible secondary mutations elsewhere in the genome, whole genome Illumina sequencing was performed commercially (SeqCenter) or in-house on at least one clone for each knockout mutant. Gene knockout at the proper genomic locus was confirmed by aligning reads from sequencing mutants to the reference wkB2 genome and knockout cassette sequences using *breseq* (v0.38.1)^99^. Mutants with successful gene deletion display missing coverage at the gene of interest and show new split-read junctions connecting the flanks of the deletion to the antibiotic resistance gene. Plasmids, gene fragments (containing deletion cassettes) and oligos (used for initial amplification of deletion cassettes and screening for successful transformants) are listed in Table S2 (Gene fragments and Oligos tabs).

pPL32 and pPL373 were cloned via Golden Gate assembly using a previously reported protocol^100^. These plasmids were initially created for another project, but used here in control strains (Fig S6). They are reported in Table S2 (Plasmids tab).

### *In vivo* bee colonization and CFU determination

To test if wkB2 WT outcompetes mutant *S. alvi* during host colonization, bees were co-colonized with 10% WT and 90% mutant. First, WT and mutant *S. alvi* grown on agar plates were resuspended in PBS and normalized to the same OD_600_ values. WT and mutant cells were then mixed at a 1:9 ratio. Resuspended cells were mixed with sucrose feed (a solution of filter-sterilized 1:1 sucrose:water^52^) at a 1:1 ratio and used to colonize approximately 20 microbiota-deficient newly emerged worker bees (NEWs) per condition, using previously described methods in which adult bees emerge in sterile containers and are then coated with the inoculant mixture, which is ingested upon grooming^100^. Bees were incubated in cup cages^100^ at 35°C and 80% relative humidity for 7-8 days. Cup cages contained pollen, which was treated with a small amount of the *S. alvi*/sucrose feed mixture on the first day. Cup cages also contained feeding tubes containing sucrose feed, which was replaced every 2-3 days, as needed. After 7 days, 11-12 bees per condition had their guts pulled on ice and ilea dissected. Ilea were resuspended in PBS, macerated with a pestle, then serially diluted in 10-fold dilutions. Serial dilutions were plated on Col-B containing Tet or Tet + Kan. Plates were incubated for 2-3 days then CFUs were counted. (Note, for plates with little to no growth, we erred on the side of over-estimation, opting to include very small colonies in our counts.) Plated bee ilea that had no detectable colonies were assigned a CFUs/ml value set at half the 20 CFUs/ml limit of detection (10 CFUs/ml), erring on the side of overestimation^101^. We then calculated CFUs/bee for all conditions and log_10_-transformed the data. A Shapiro-Wilk test was performed on log_10_-transformed data to test for normality. Individual data points and group medians with 95% CI were plotted. Significant differences between the indicated log_10_-transformed group medians were determined by Mann-Whitney U tests with the two-stage Benjamini, Krieger, & Yekutieli method to correct for multiple comparisons by controlling the FDR. N = 11-12 biological replicates for each condition.

To assess whether mutant *S. alvi* strains can mono-colonize bees, a workflow similar to the co-colonization experiment was employed, utilizing either 100% mutant (experimental group) or 100% WT (control group). Independent mono-colonization experiments with several of the same strains were conducted to confirm the results for those strains (Fig. 1B-C, Fig. S4). First, plate-grown WT and mutant *S. alvi* cells were resuspended in PBS. Cells were then normalized to the same OD_600_ for Fig. 1C. OD_600_ normalization was not used in all experiments, however, as inoculant concentration is not a limiting factor in *S. alvi* colonization down to 50 CFUs/bee^102^. Resuspended cells were subsequently mixed with sucrose feed at a 1:1 ratio (Fig. 1B-C) or 4:1 ratio (Fig. S4) and used to coat approximately 20 microbiota-deficient NEWs (newly emerged bees) per condition. Inoculated bees were incubated for 7 days (Fig. 1C; Fig S4) or 8 days (Fig. 1B). Ileum dissection, CFU plating, and CFU analysis were then performed, following the same workflow used for co-colonized bees. CFUs/bee were calculated for each condition and log_10_-transformed. A Shapiro-Wilk test was performed on log_10_-transformed data to test for normality. Individual data points and group medians with 95% CI were plotted. For each experiment, Mann-Whitney U tests (with the two-stage Benjamini, Krieger, & Yekutieli method to correct for multiple comparisons by controlling the FDR) were performed to determine significant differences between the log_10_-transformed medians of WT (+Tet) and all other groups. N = 7-12 (Fig. 1B); N = 12 (Fig. 1C); N = 4-6 (Fig. S4) biological replicates for each condition. Note that a single experiment was conducted to produce the data shown in both Fig. 1A and Fig. 1C. Therefore, the WT (+Tet and +Tet + Kan) data are identical across these panels. The graphs were split to improve clarity.

### *In vitro* bacterial fitness determination

*S. alvi* fitness was initially assessed by *in vitro* CFU determination. Briefly, *S. alvi* WT and mutant strains were grown in liquid culture for 3 days and OD_600_ normalized. OD_600_-normalized cells were diluted in a 10-fold dilution series, plated on Col-B agar, and incubated for 2-3 days. After 2-3 days, CFUs were counted and CFUs/ml were calculated. A Shapiro-Wilk test of untransformed data was performed to test for normality. Data were then log_10_-transformed. Individual data points and group medians with 95% CI of log_10_ transformed data were then plotted for each condition. Significant differences between the log_10_-transformed group medians were assessed by performing a Kruskal–Wallis test with Dunn’s multiple comparisons.

*S. alvi* fitness was additionally assessed by growth curve analysis. Briefly, *S. alvi* wkB2 WT and mutant strains were grown in liquid culture for 3 days. Cells were diluted to an OD_600_ of 0.025 (Fig S5B) or 0.125 (Fig S5C-E) in 96-well plates containing Columbia broth or Insectagro, as indicated. 96-well plates were incubated in a Spark 10M plate reader (Tecan, Switzerland) for 48 or 72 hours at 35°C, 5% CO_2_. OD_600_ readings were taken every hour, with 30 s of shaking prior to each OD_600_ reading. Growth curve data was plotted for each condition. Each trendline represents a single biological replicate. For Figure S5B, N=1 biological replicate (with 3 technical replicates per condition). For Figure S5C-E, N=3 biological replicate (with 3 technical replicates per biological replicate). Trendline = mean of technical replicates; error bars = SD.

### *In vitro* biofilm production assay

*In vitro* biofilm production was assessed using the crystal violet biofilm assay^51–53^. *S. alvi* wkB2 WT and mutant strains were grown in liquid culture for 3 days. Any biofilm present was resuspended and 96-well plates were inoculated with a 1:40 dilution of resuspended cells. Unless otherwise indicated, cells were grown in Insectagro. (*S. alvi* was grown in BHI in two experiments, shown in figure S7). Outer wells were filled with media or culture to prevent evaporation of inner wells. These wells were not included in experimental results. Plates were incubated for 2 days. OD_600_ was then measured for plates using a Spark 10M plate reader. Plates were subsequently washed with dH_2_O and stained with 0.1% w/v crystal violet in ddH_2_O. Wells were washed to remove unbound crystal violet and stained plates were imaged. 30% v/v acetic acid in ddH_2_O was added to wells to resuspend any remaining crystal violet and OD_550_ was measured for plates to quantify the amount of crystal violet present, itself a proxy for biofilm production. Biofilm formation was normalized to cell density by dividing OD_550_ by OD_600_. Absorbance (OD_550_, OD_600_, and OD_550_/OD_600_) was plotted for each condition. Shapiro-Wilk tests were performed to test for normality. Group medians with individual data points and 95% CI were graphically displayed. Significance between the indicated groups was determined by performing Mann-Whitney U tests or a Kruskal-Wallis test (specified in figure legends) with the two-stage Benjamini, Krieger, & Yekutieli method to correct for multiple comparisons by controlling the FDR.

### Cell sedimentation assay

*S. alvi* cell sedimentation was assessed by measuring the rate at which cells settle in culture tubes, using previously described protocols^74,75^. Briefly, WT and Δ*staA* cells were grown in liquid culture for 3 days, to allow for WT to form biofilm. After 3 days, cells were photographed, resuspended, and OD_600_-normalized. Cells were then transferred to 15 ml conical tubes and photographed again. Following resuspension, cells were allowed to settle for 120 minutes. At various timepoints, more frequent at the beginning of the experiment, a small volume of culture was sampled from the top of each tube and transferred to a 96-well plate, which was kept on ice to prevent cell growth. After the final timepoint, the 15 ml culture tubes were again photographed. OD_600_ was also measured for the 96-well plate containing culture samples using a Spark 10M plate reader. Absorbance data were normalized to the initial OD_600_ value for each condition. Normalized datapoints (means of replicates) were then plotted with error bars (SD) on a log-scale. For each condition, a non-linear best-fit trendline was generated using a two-phase decay model. Trendlines were added to the plot. N= 3 biological replicates for each condition.

### Light microscopy

Auto-aggregation of *S. alvi* was visualized by light microscopy. Briefly, WT and Δ*staA* cells were grown in liquid culture for 3 days to allow for WT to form biofilm. After 3 days, WT cells (present in biofilm) were dislodged from the walls of the culture tube by scraping with a sterile inoculating loop, then rubbing the loop onto a slide before covering them with a cover slip for imaging. Δ*staA* cells (which are planktonic) were directly deposited in liquid culture onto a slide for imaging. Mounted cells were then imaged on a Zeiss Axiovert 200M microscope with a 63X objective under differential interference contrast (DIC). Images were processed for publication in Fiji^103^.

### Flow cytometry

Auto-aggregation of *S. alvi* was further assessed using flow cytometry, as previously described in other bacteria^104^. Briefly, WT and Δ*staA* colonies were scraped from agar plates, resuspended in PBS, then standardized to an OD of 0.1. Samples were subsequently loaded onto the stage of a LSRFortessa (BD Biosciences, San Jose, CA). Instrument settings were adjusted to record cell size (FSC/SSC), and flow cytometric analysis was then performed for each sample. Data were collected using FACSDiva (BD Biosciences, USA) software and analyzed using FlowJo (BD Biosciences, USA). For each sample, the histogram of the number of events with different FSC-H values was graphed to assess and compare auto-aggregation.

### Scanning electron microscopy

WT and mutant *S. alvi* cells were visualized by SEM. First, WT and Δ*staA* cells were grown for three days in liquid culture, to allow for biofilm formation by WT. After 3 days, planktonic and biofilm cells were resuspended, then added to Aclar or glass substrates. Samples were dried, then fixed with 2% glutaraldehyde + 0.15% ruthenium red for 1 hour, then stained with 0.15% ruthenium red overnight. The next day, samples were washed with 0.1M sodium cacodylate buffer, post-fix stained with 1% osmium tetroxide + 0.15% ruthenium red for 2 hours, then washed in water overnight. The next day, samples were dehydrated using an ethanol/HMDS gradient, then dried in air overnight. The following day, stained substrates were mounted to stubs with conductive paint and sputter coated with platinum/palladium to a thickness of 12 nm. Samples were imaged on a Zeiss Supra 40V Scanning Electron Microscope. Scale bars and annotations were added to micrographs in Microsoft PowerPoint.

### Protein structure prediction

Protein structures were predicted for TAAs from *S. alvi* and other bacteria using AlphaFold^105^. AlphaFold prediction fidelity decreases as protein size increases, so TAA structures were predicted using an iterative, manual segmentation process. Accordingly, protein sequences were initially segmented into ∼400-800 amino acid chunks. The signal sequence was omitted from structure prediction, as this peptide would be cleaved during TAA export from the inner membrane.

A first round of structure prediction was performed by uploading the segmented TAA sequences to Tamarind^106^, a webserver that employs AlphaFold for structure prediction. Since TAAs form trimers, structure prediction was performed in multimer mode using three identical sequences of each TAA segment. Initial partial structures were generated and then examined using Tamarind’s protein viewer to identify any of the following issues: 1) A low fold score, 2) Clear folding errors (such as disordered regions in areas expected to form alpha helices), or 3) Significant dissimilarity to TAA motifs, domains, or full structures found in other species^55,84,107^. Flawed structures were further divided into smaller segments, re-uploaded to Tamarind, and re-checked for issues in an iterative process until flaws were sufficiently mitigated. Adjacent segments were designed to overlap by 50-100 residues, to aid in eventual concatenation of segments. For all domains but one, quality structures were obtained for each TAA after multiple rounds of segmentation and structure prediction. The lone domain that did not fold was the unassigned domain, an unstructured region C-terminal to the signal sequence and N-terminal to the beta helix head. As such, the unassigned domain was omitted from final structures.

Domain structures considered to be of high quality (i.e., free from obvious defects) were downloaded and sequentially concatenated in PyMOL^108^. Overlapping residues were then removed and monomers, loops, and/or residues were color coded. Cartoons with or without side chains, and surface exposed structures were rendered with ray tracing. In select figures, side chains were labeled. The length of each TAA was also measured in PyMOL. Structures are depicted on the same scale in Fig 5B. Visual presentation of *S. alvi* TAA structures was modeled after figure panels in Bassler et al 2015^84^.

### Creation of protein domain diagrams

Protein domain diagrams were created for TAAs from *S. alvi* and other species, using previously reported TAA domain diagrams as templates^89^. Briefly, TAA domain annotations in UniProt^109^ were used to determine the lengths of the signal peptides. The lengths of subsequent domains were determined by manual assessment of domain boundaries in each predicted protein structure. Unstructured residues C-terminal to the signal sequence and N-terminal to the canonical beta helix head region were classified as unassigned. Finally, cartoon diagrams of each protein were drawn with labeled domains. Protein domain diagrams were drawn to the same scale (diagram length/# of amino acids).

### Phylogenetic analysis of TAAs across *Snodgrassella*

To construct a TAA phylogeny, a list of StaA, StaB, and StaC orthologs within *Snodgrassella* was generated. To do this, BLASTP searches were performed on NCBI using the wkB2 StaA, StaB, or StaC anchor domains as inputs. Outputs with lower identity to the input were cross-referenced in searches for the other paralogs, to determine if an output had highest percent identity to StaA, StaB, or StaC. Orthologs with partial (i.e., incomplete) protein sequences in NCBI were omitted from our ortholog list. Additionally, only orthologs present in the *Snodgrassella* species highlighted in Cornet et al 2022^61^ were included in this analysis. A FASTA file was then generated containing the StaA ortholog anchor sequences and wkB2 StaB and StaC anchor sequences as outgroups. Using this FASTA file as an input, a multiple sequence alignment (MSA) of the StaA ortholog anchors and StaB/C outgroups was generated using Clustal Omega^110^. The output Newick file was saved and any small negative values were manually changed to “0”. Using this Newick file as an input, a phylogenetic tree rooted in StaB/C was then created in iTOL (v6)^111^. The tree was resized and children were rotated to cluster taxonomic groups in the order presented in Cornet et al 2022^61^. Taxonomic groups were then color-coded and labeled using previous designations^61^ (A1-2: *Apis*-specific *Snodgrassella*; B2-5: *Bombus*-specific *Snodgrassella*). Finally, our list of StaB/C orthologs was manually searched for each species on the tree. Presence or absence of StaB/C orthologs in each species was then recorded and indicated next to the tree.

### Phylogenetic and residue conservation analysis of StaA neck+stalk repeats

Phylogenetic and residue conservation analyses were performed for StaA neck+stalk repeats. First, a list of wkB2 StaA neck+stalk repeat sequences was generated. The predicted protein structure of StaA was used to visually pinpoint the beginning and end of each neck+stalk repeat. The sequence for the full length of each repeat was then recorded. Next, an MSA of all 27 StaA neck+stalk protein sequences was performed using Clustal Omega^110^. Clustal Omega-generated figures, including a phylogenetic tree and an alignment with color-coded residues, were then downloaded and combined into a single figure panel.

A sequence logo illustrating residue conservation among the StaA neck+stalk repeats was subsequently generated using WebLogo (version 2.8.2)^112^. A file of the StaA repeat alignment in FASTA format converted by Seqret was downloaded from Clustal Omega and uploaded to the WebLogo server. Sequence logos for the neck and two halves of the stalk were downloaded and annotated.

### Conserved residue analysis of TAA signal sequences in *Snodgrassella*

StaA and StaB protein sequences were assessed for conservation across *Snodgrassella*. To do this, a multiple sequence alignment (MSA) of *Snodgrassella* StaA ortholog protein sequences was performed using Clustal Omega^110^. A separate MSA of StaB orthologs was performed. Alignment files were then uploaded to the ConSurf web server^113,114^, using wkB2 StaA/StaB as the query sequences. In ConSurf analysis, each residue is assessed for degree of conservation across the input sequences, then color coded accordingly. Conserved residue output files were downloaded, then assessed for conservation within different domains. ConSurf alignments of the highly conserved StaA/StaB signal sequences were reported here. Consensuses for the signal sequences were determined and also reported.

### Multiple sequence alignment between StaA and StaB head domains

Sequence variability and conservation between the StaA and StaB head domains were assessed. To do this, an MSA of the wkB2 StaA and StaB head domains was performed using Clustal Omega^110^. An output file was obtained that indicated if residues were identical, conservatively mutated, or semi-conservatively mutated. A region of low identity was identified and highlighted in each ortholog with a color specific to that ortholog. Charged residues in this region were also color coded to indicate sign of charge.

### Antimicrobial susceptibility assay

Biofilm-mediated protection from antimicrobials was tested by exposing antimicrobials to 2-day old liquid cultures. Briefly, a 96-well plate was inoculated with 3-day old overnights of WT and Δ*staA* to an OD_600_ of 0.025 in Insectagro. Plates were incubated for 2 days to allow biofilm to form. After 12 days, gentamicin or apidaecin 1B (NovoPro Bioscience Inc., China) were added to the indicated concentrations in a 2-fold dilution series. Plates were incubated with antimicrobials overnight to allow for cell killing. The next day, plates were centrifuged for 12 minutes at 2038 x *g* to pellet any planktonic cells, and media was aspirated to remove excess antimicrobial. Cells were scraped and resuspended in PBS. Resuspended cells were then diluted in a 10-fold dilution series, plated on Col-B agar plates, and incubated for 2-3 days. After 2-3 days, CFUs were counted and CFUs/ml were calculated. Antimicrobial concentrations that resulted in no detectable colonies were assigned a CFUs/ml value set at half the 200 CFUs/ml limit of detection (100 CFUs/ml), which errs on the side of overestimation^101^. Fold change in CFUs/ml was determined by normalizing values to the CFUs/ml values for cells not exposed to antimicrobials. Normalized data were then log_10_-transformed. For each group, individual data points and medians with 95% CI were subsequently plotted. N = 1-3 (Fig 6A) or N = 2 (Fig 6B) biological replicates per condition.

## Acknowledgements

We thank members of the Moran and Barrick labs for helpful discussions. We thank Kim Hammond and Eli Powell for providing apiculture training and bee hive maintenance. We thank Dr. Daniel Deatherage for performing whole genome sequencing to confirm gene knockout in engineered mutants. We thank Dr. Alan Emanuel Silva Cerqueira for providing advice on phylogenetic tree construction. We thank Tyler de Jong for helping create plasmids pPL32 and pPL373. We also thank Michelle Mikesh for assistance in performing SEM. Flow cytometry was performed at the Center for Biomedical Research Support Microscopy and Imaging Facility at UT Austin (RRID:SCR_021756). Figures were made using BioRender.

This work was supported by funding from the USDA NIFA (2023-67012-39356 to P.J.L.), the National Science Foundation (IOS-2103208 to J.E.B. and N.A.M.), the U.S. Army Research Office (W911NF-20-1-0195 to J.E.B. and N.A.M.), the National Institutes of Health (R35GM131738 to N.A.M.), and the UT Austin College of Natural Sciences (Spark Grant to J.E.B).

## Author contributions

Conceptualization, P.J.L., N.A.M., and J.E.B.; Methodology, P.J.L., A.Z.A., and I.G.; Investigation, P.J.L., A.Z.A., I.G., and S.L.T.; Writing, P.J.L., A.Z.A., and I.G.; Editing, P.J.L., A.Z.A., I.G., S.L.T., N.A.M., and J.E.B.

## Competing interests

N.A.M. and J.E.B. have a patent (US11382989B2) on the use of engineered symbionts to improve bee health.

## Supplemental material

**Table S1. StaA/StaB length across *Snodgrassella* (see excel doc)**

**Table S2. Strains, plasmids, gene fragments, and oligos (see excel doc)**

## Supplemental Figures

**Figure S1.**
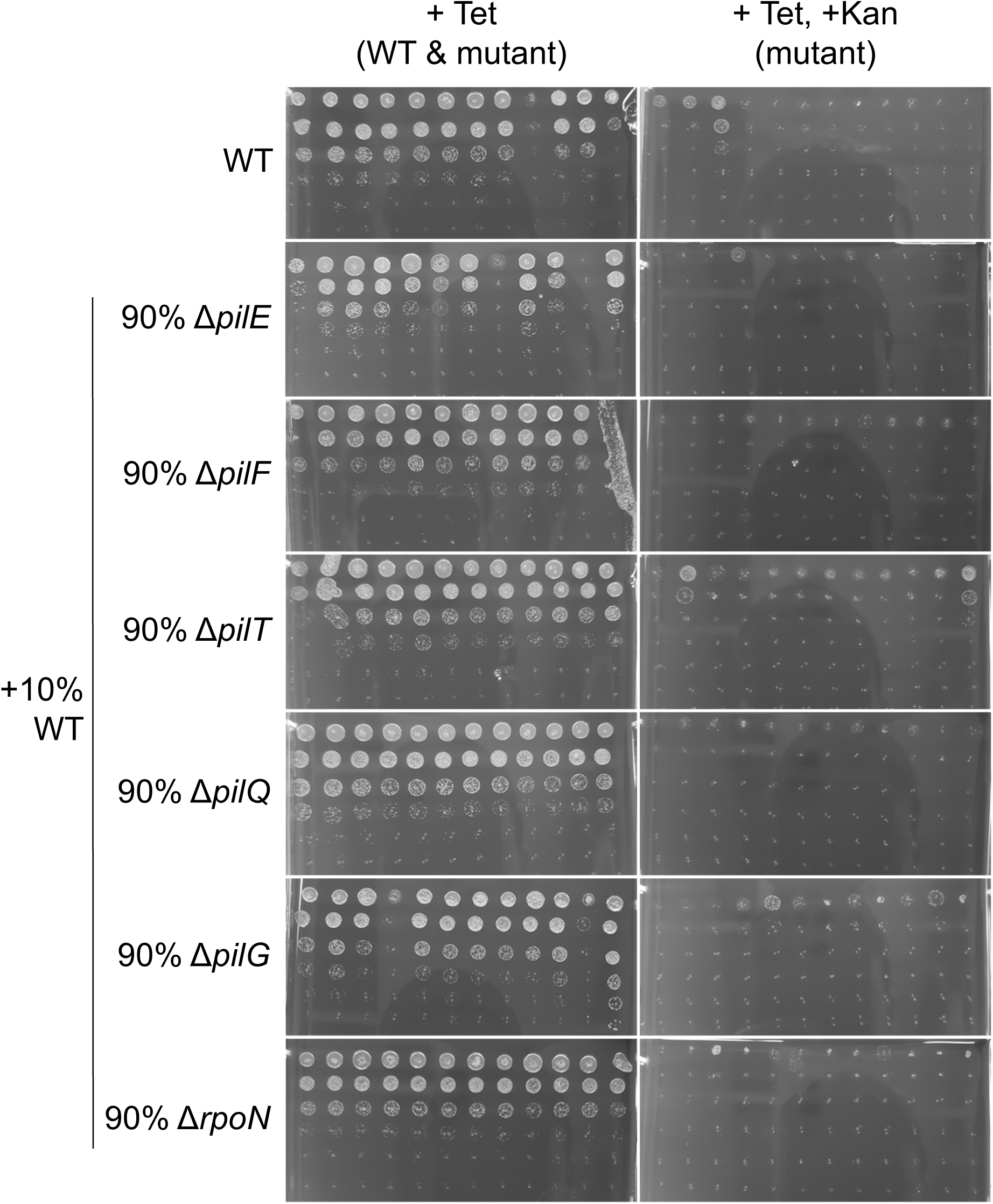
Viability of WT and mutant *S. alvi* cells co-colonized in bees. CFU plates of WT and mutant *S. alvi* cells co-colonized in bees used to generate CFU plots in Figure 1A. Ilea from bees co-colonized with 10% WT and 90% mutant *S. alvi* were plated on media containing Tet (selects for WT and mutant) and Tet, Kan (selects for mutant). N = 12 for each condition. Each column is a 10-fold dilution series of a macerated ileum. Each column is an individual bee replicate.

**Figure S2.**
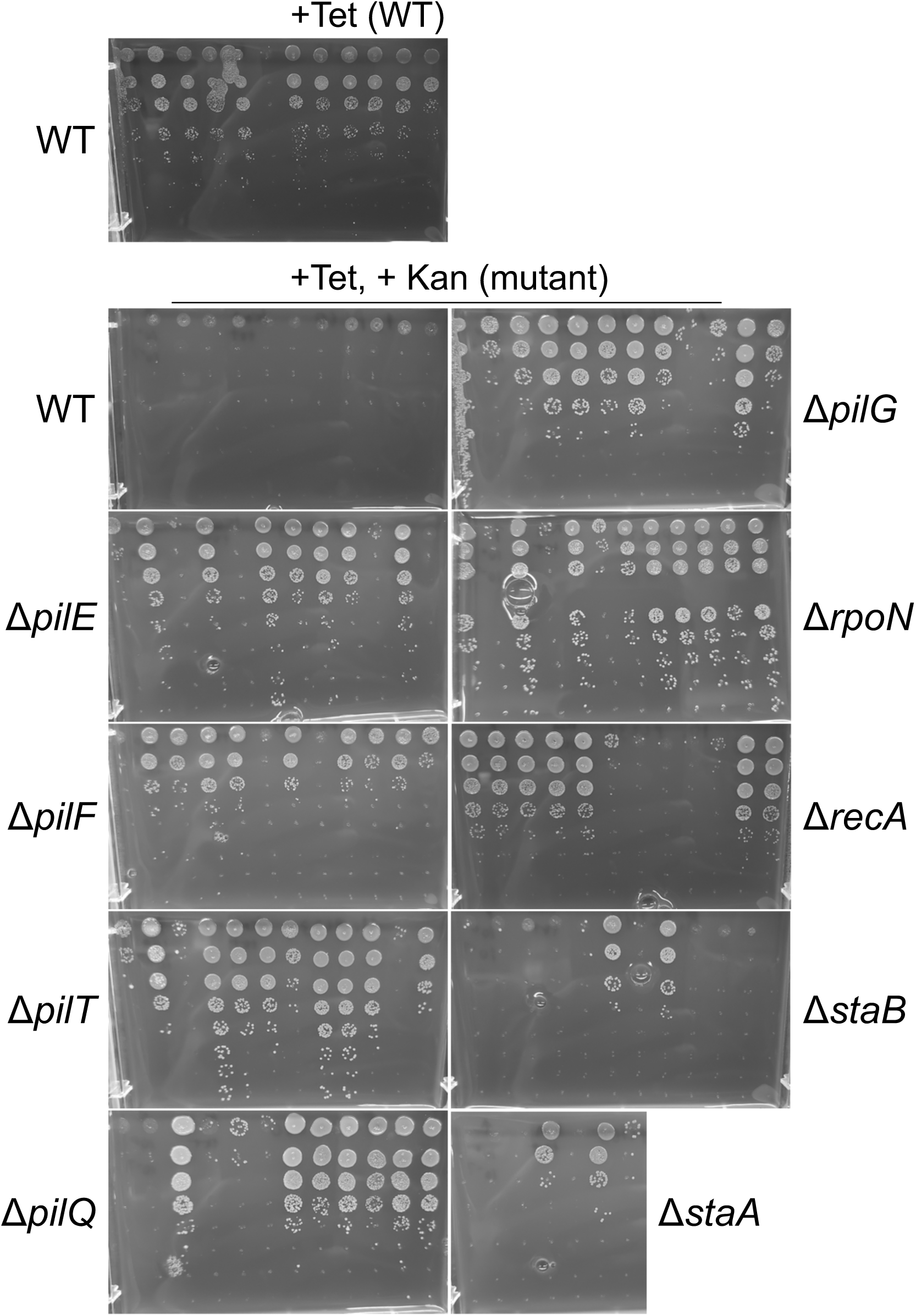
Viability of mutant *S. alvi* cells mono-colonized in bees (experiment 1) CFU plates of WT and mutant *S. alvi* cells mono-colonized in bees used to generate CFU plots in Figure 1B (experiment 1). Ilea from bees mono-colonized with mutant *S. alvi* were plated on media containing Tet, Kan (selects for mutant). Ilea from bees mono-colonized with WT control were plated on both Tet (allows for WT growth) and Tet, Kan (does not allow for WT growth). N = 7-12 for each condition. Each column is a 10-fold dilution series of a macerated ileum. Each column is an individual bee replicate.

**Figure S3.**
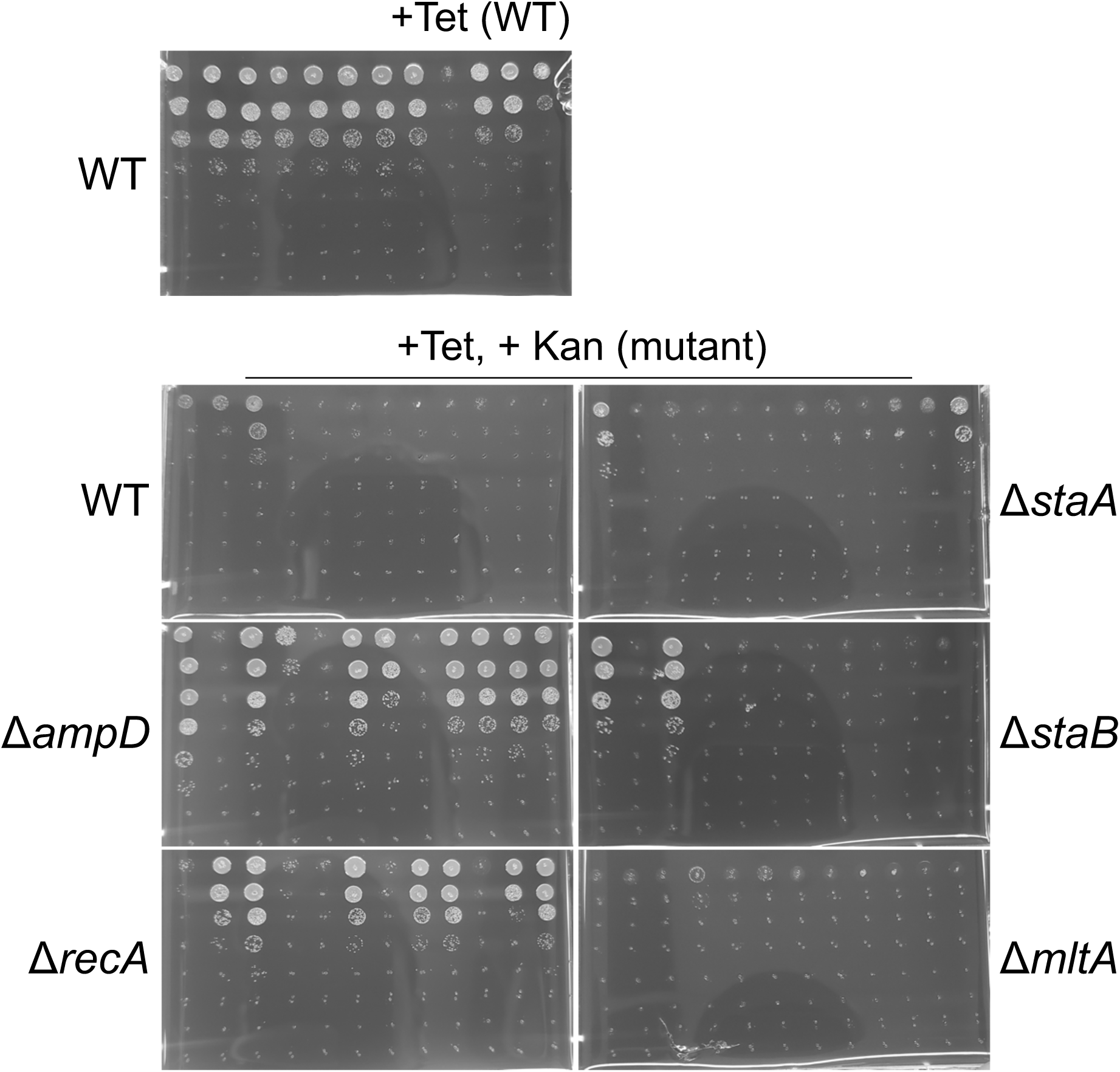
Viability of mutant *S. alvi* cells mono-colonized in bees (experiment 2) CFU plates of WT and mutant *S. alvi* cells mono-colonized in bees used to generate CFU plots in Figure 1C (experiment 2). Ilea from bees mono-colonized with mutant *S. alvi* were plated on media containing Tet, Kan (selects for mutant). Ilea from bees mono-colonized with WT control were plated on both Tet (allows for WT growth) and Tet, Kan (does not allow for WT growth). N = 12 for each condition. Each column is a 10-fold dilution series of a macerated ileum. Each column is an individual bee replicate.

**Figure S4.**
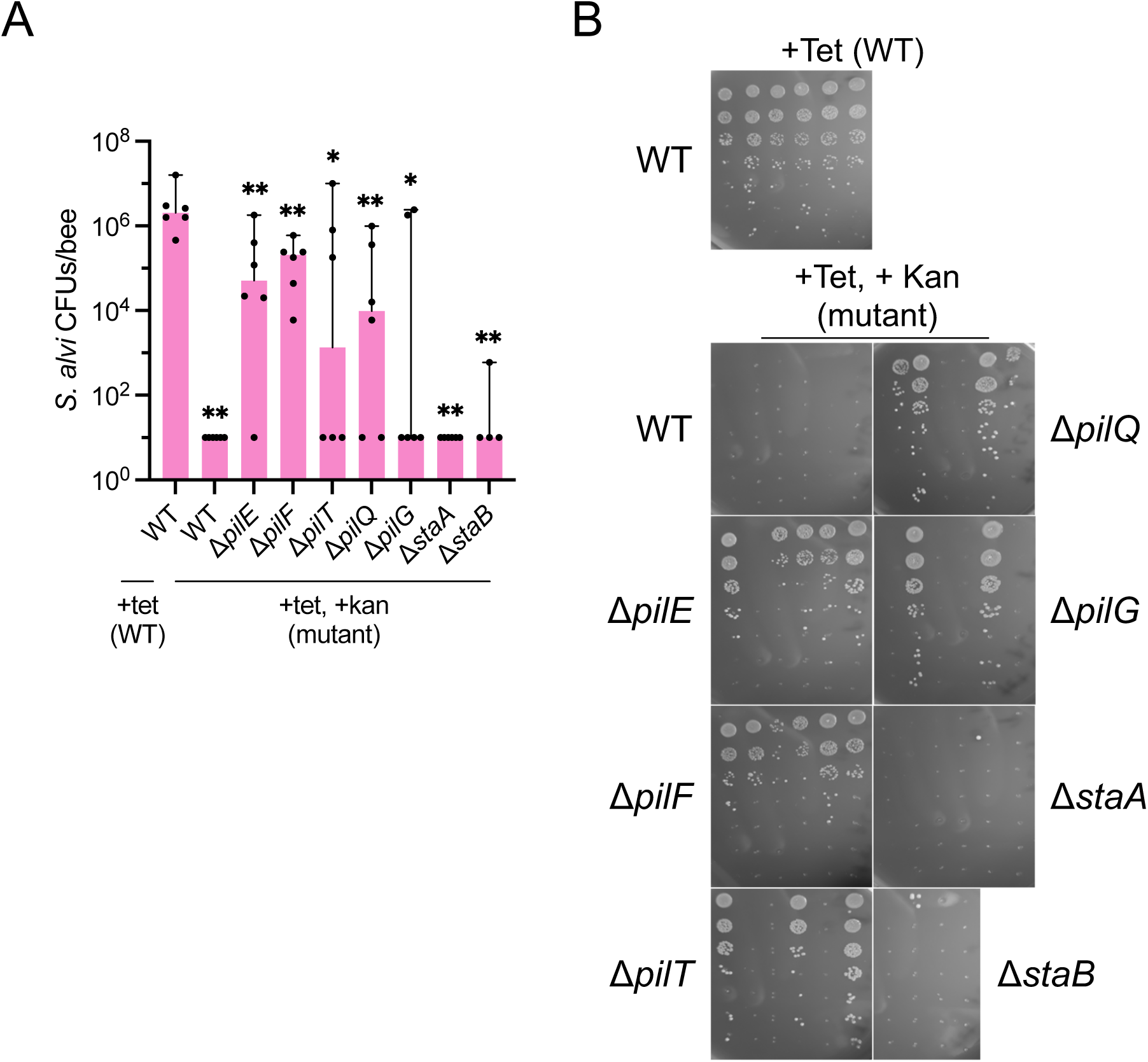
Viability of mutant *S. alvi* cells mono-colonized in bees (experiment 3) A. Plot of CFU counts of *S. alvi* mutants mono-colonized in bees, confirming that Δ*staA* and Δ*staB* cannot effectively mono-colonize bees. Ilea from bees mono-colonized with mutant *S. alvi* were plated on media containing Tet, Kan (selects for mutant). Ilea from bees mono-colonized with WT control were plated on both Tet (allows for WT growth) and Tet, Kan (does not allow for WT growth). N = 4-6 biological replicates for each condition. A Shapiro-Wilk test found that not all log_10_-transformed data are normally distributed. Individual data points and group medians with 95% CI are shown. Significant differences between the log_10_-transformed medians of wkB2 (+Tet) and all other groups were determined by Mann-Whitney U tests with the two-stage Benjamini, Krieger, & Yekutieli method to correct for multiple comparisons by controlling the FDR (\**q* ≤ 0.05, \*\**q* ≤ 0.01). B. CFU plates of WT and mutant *S. alvi* cells mono-colonized in bees used to generate CFU plots in Figure S4A (experiment 3). Ilea from bees mono-colonized with mutant *S. alvi* were plated on media containing Tet, Kan (selects for mutant). Ilea from bees mono-colonized with WT control were plated on both Tet (allows for WT growth) and Tet, Kan (does not allow for WT growth). N = 4-6 for each condition. Each column is a 10-fold dilution series of a macerated ileum. Each column is an individual bee replicate.

**Figure S5.**
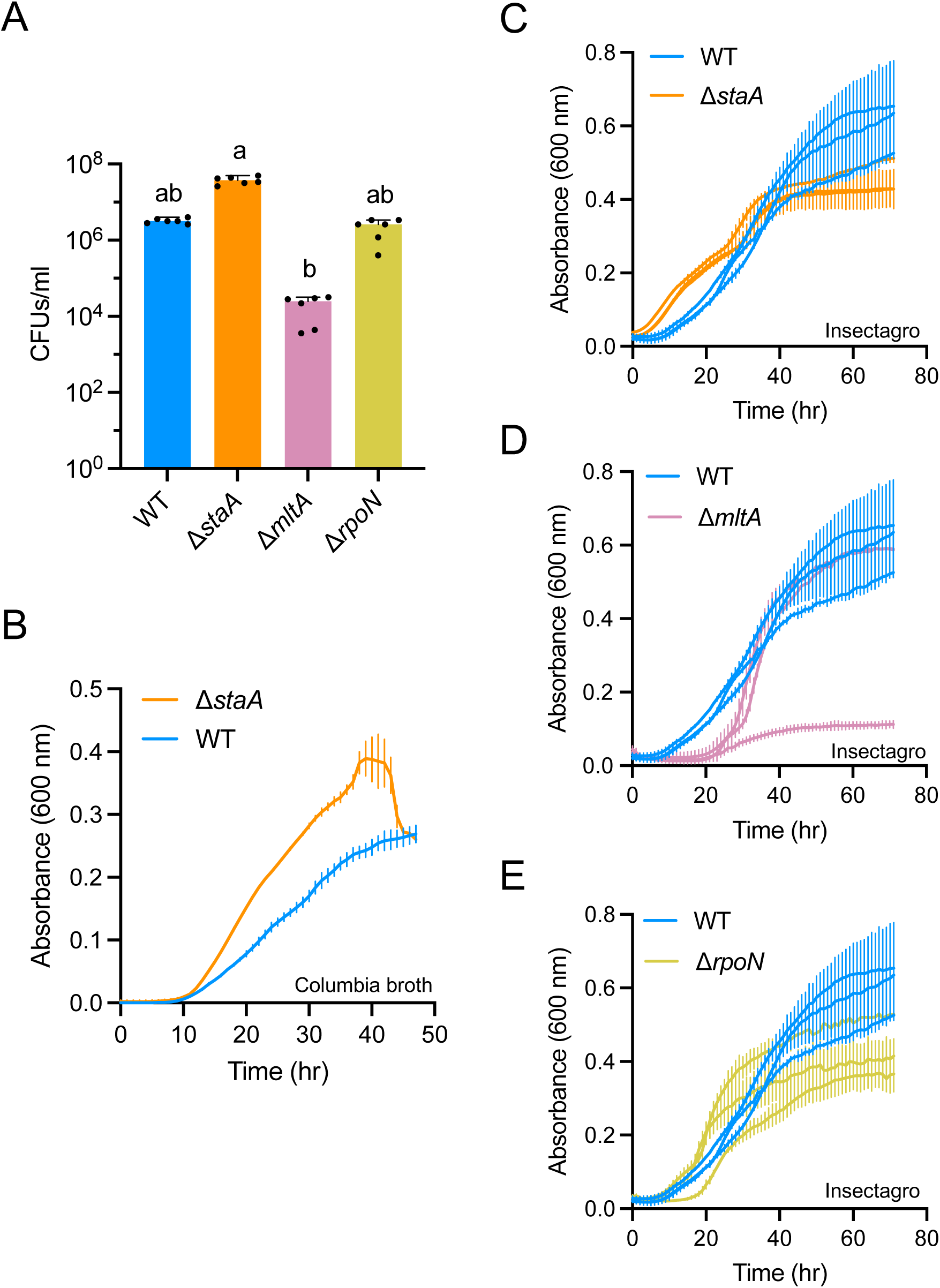
Fitness of Δ*staA*, Δ*mltA,* and Δ*rpoN* cells is not substantially reduced compared to wkB2. A. Plot of CFU counts of *S. alvi* mutants grown *in vitro*, indicating Δ*staA* and Δ*rpoN* has WT-like or better viability and Δ*mltA* has slightly reduced viability. N = 6 biological replicates for each condition. A Shapiro-Wilk test of untransformed data found that not all data are normally distributed. Individual datapoints and group medians with 95% CI of log_10_ transformed data are shown. Significant differences between log_10_-transformed group medians (P < 0.0001; found by a Kruskal–Wallis test with Dunn’s multiple comparisons) are indicated by dissimilar letters above groups. B. Plot of growth curve of Δ*staA* and wkB2 grown in Columbia broth, indicating Δ*staA* grows better than wkB2 in Columbia broth. Single biological replicates were run for each condition. Each biological replicate was run in technical triplicate. Trendline = mean of technical replicates; error bars = SD. C. Plot of growth curve of Δ*staA* and wkB2 grown in Insectagro, indicating Δ*staA* growth is slightly reduced compared to wkB2 in Insectagro. N = 3 biological replicates for each condition. Each biological replicate was run in technical triplicate. Trendline = mean of technical replicates for each biological replicate; error bars = SD. D. Plot of growth curve of Δ*mltA* and wkB2 grown in Insectagro, indicating Δ*mltA* growth is comparable to wkB2 in Insectagro in 2/3 replicates. N = 3 biological replicates for each condition. Each biological replicate was run in technical triplicate. Trendline = mean of technical replicates for each biological replicate; error bars = SD. E. Plot of growth curve of Δ*rpoN* and wkB2 grown in Insectagro, indicating Δ*rpoN* growth is slightly reduced compared to wkB2 in Insectagro. N = 3 biological replicates for each condition. Each biological replicate was run in technical triplicate. Trendline = mean of technical replicates for each biological replicate; error bars = SD.

**Figure S6.**
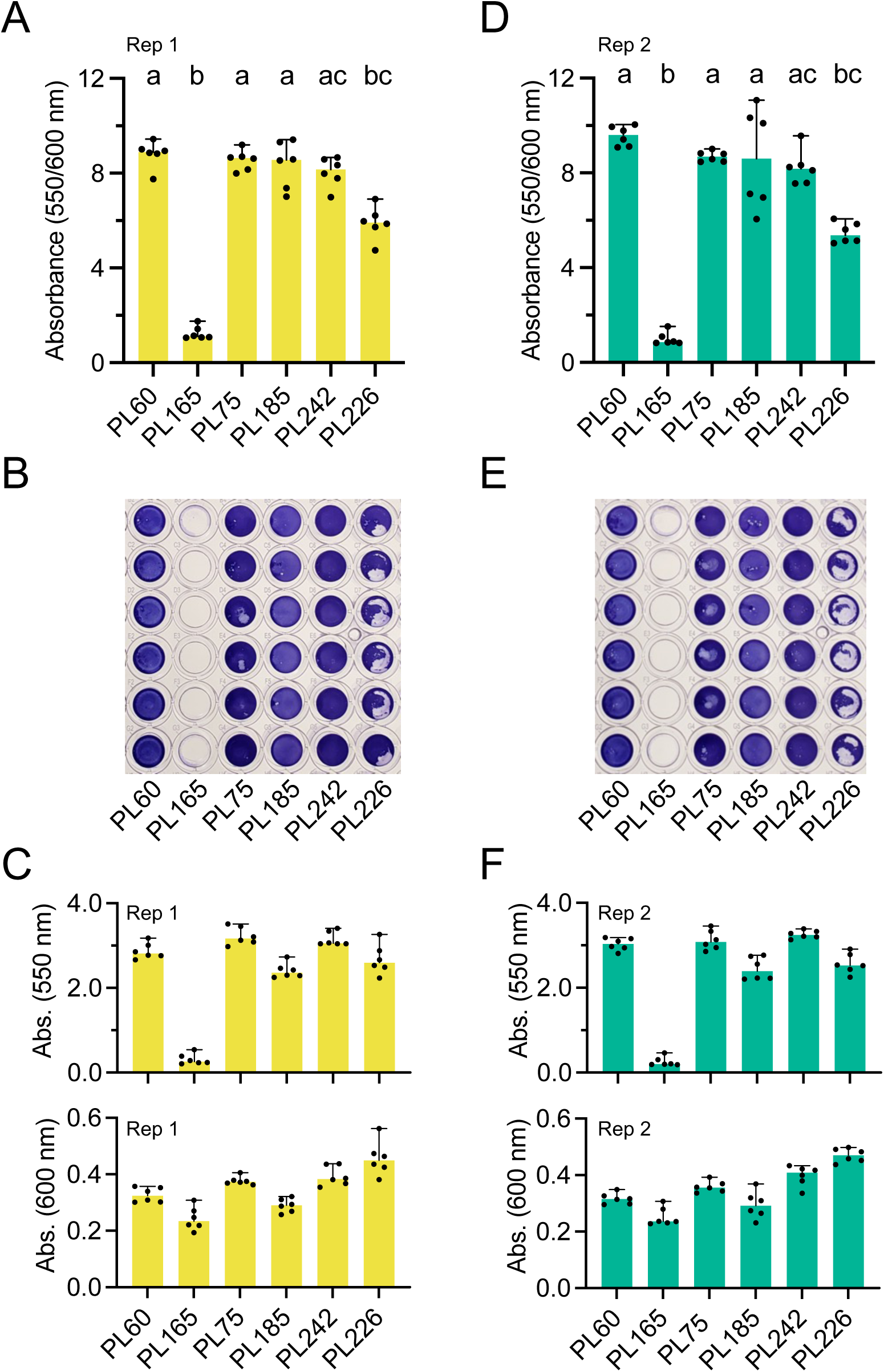
Biofilm formation is not affected by mutation in SALWKB2_RS07515. A. Plot quantifying cell growth-normalized biofilm formation, indicating mutation in SALWKB2_RS07515 (present in PL75) does not alter biofilm formation (rep 1). N = 6 for each condition. A Shapiro-Wilk test found that not all data are normally distributed. Individual data points and group medians with 95% CI are shown. Significant differences between group medians (*q* < 0.05; found by a Kruskal–Wallis test with the two-stage Benjamini, Krieger, & Yekutieli method to correct for multiple comparisons by controlling the FDR) are indicated by dissimilar letters above groups. B. Image of 96-well plate stained with crystal violet, indicating mutation in SALWKB2_RS07515 (present in PL75) does not alter biofilm formation (rep 1). C. Plots quantifying non-normalized biofilm formation (top) and cell growth (bottom), indicating PL75 (mutation in SALWKB2_RS07515) has WT-like growth and biofilm formation (rep 1 N = 6 for each condition. A Shapiro-Wilk test found that not all data are normally distributed. Individual data points and group medians with 95% CI are shown. D. Plot quantifying cell growth-normalized biofilm formation, indicating mutation in SALWKB2_RS07515 (present in PL75) does not alter biofilm formation (rep 2). N = 6 for each condition. A Shapiro-Wilk test found that not all data are normally distributed. Individual data points and group medians with 95% CI are shown. Significant differences between group medians (*q* < 0.05; found by a Kruskal–Wallis test with the two-stage Benjamini, Krieger, & Yekutieli method to correct for multiple comparisons by controlling the FDR) are indicated by dissimilar letters above groups. E. Image of 96-well plate stained with crystal violet, indicating mutation in SALWKB2_RS07515 (present in PL75) does not alter biofilm formation (rep 2). F. Plots quantifying non-normalized biofilm formation (top) and cell growth (bottom), indicating PL75 (mutation in SALWKB2_RS07515) has WT-like growth and biofilm formation (rep 2). N = 6 for each condition. A Shapiro-Wilk test found that not all data are normally distributed. Individual data points and group medians with 95% CI are shown.

**Figure S7.**
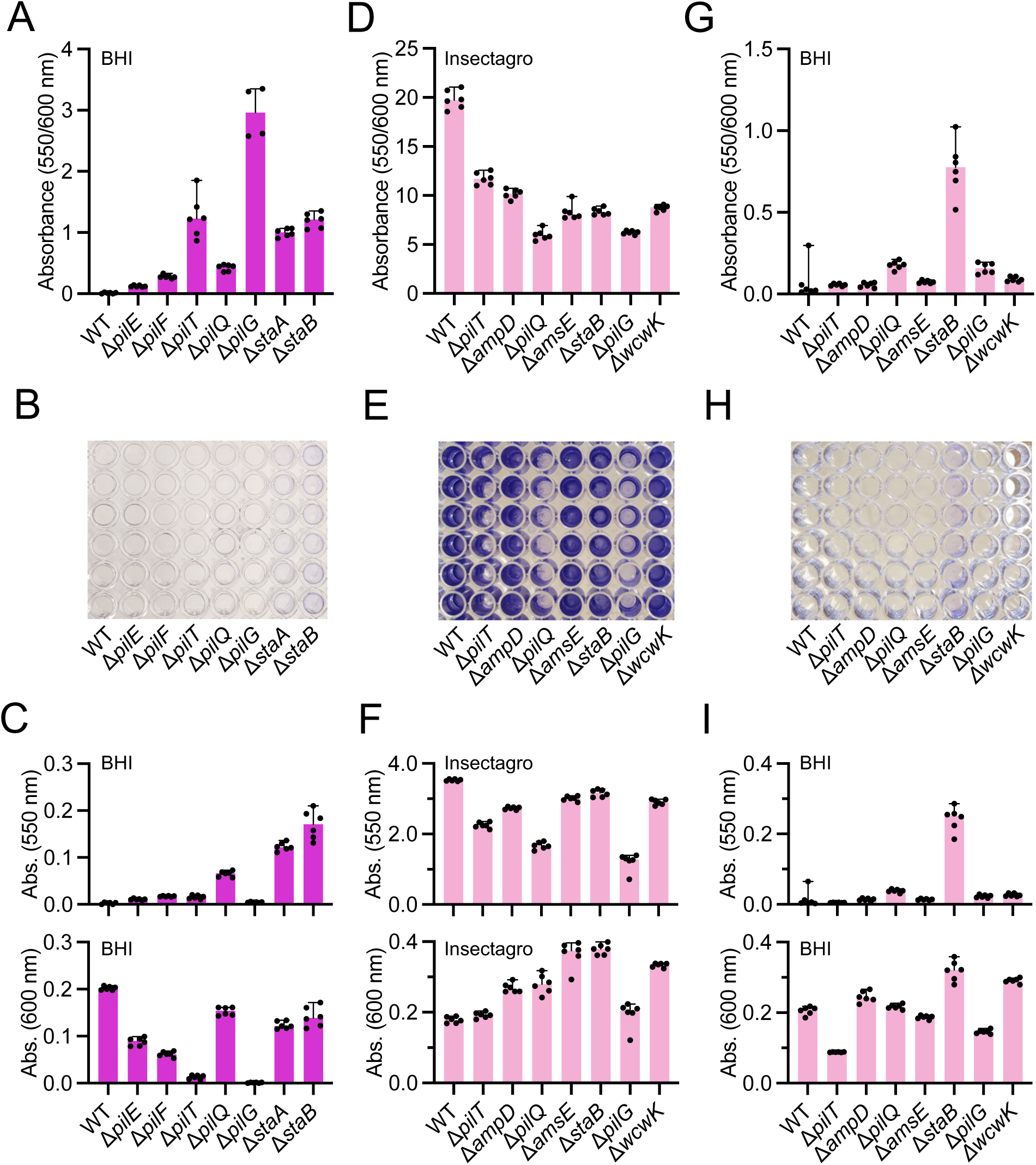
Ability of *S. alvi* to form biofilm *in vitro* is media-dependent. A. Plot quantifying cell growth-normalized biofilm formation in BHI, indicating WT and adhesin *S. alvi* mutants cannot form biofilm in BHI. N = 6 for each condition except for Δ*pilG*, where N = 4. Individual data points and group medians with 95% CI are shown. (Data were found to be normally distributed by Shapiro-Wilk test, but group medians are shown to maintain consistency across the figure). B. Image of 96-well plate stained with crystal violet, indicating WT and adhesin mutant *S. alvi* strains do not form biofilm in BHI. C. Plots quantifying non-normalized biofilm formation (top) and cell growth (bottom), indicating WT and most adhesin mutants grow in BHI, but do not form biofilm. N = 6 for each condition. A Shapiro-Wilk test found that not all data are normally distributed. Individual data points and group medians with 95% CI are shown. D. Plot quantifying cell growth-normalized biofilm formation in Insectagro, indicating WT and some *S. alvi* mutants can form robust biofilms in Insectagro. N = 6 for each condition. A Shapiro-Wilk test found that not all data are normally distributed. Individual data points and group medians with 95% CI are shown. E. Image of 96-well plate stained with crystal violet, indicating WT and some mutant *S. alvi* strains do form robust biofilms in Insectagro. F. Plots quantifying non-normalized biofilm formation (top) and cell growth (bottom). All strains grow in Insectagro and some strains, including WT, form robust biofilms in Insectagro. N = 6 for each condition. A Shapiro-Wilk test found that not all data are normally distributed. Individual data points and group medians with 95% CI are shown. G. Plot quantifying cell growth-normalized biofilm formation in BHI, indicating WT and additional *S. alvi* mutants cannot form biofilm in BHI. N = 6 for each condition. A Shapiro-Wilk test found that not all data are normally distributed. Individual data points and group medians with 95% CI are shown. H. Image of 96-well plate stained with crystal violet, indicating WT and additional mutant *S. alvi* strains do not form biofilm in BHI. I. Plots quantifying non-normalized biofilm formation (top) and cell growth (bottom), indicating WT and most additional mutants grow in BHI, but do not form biofilm. N = 6 for each condition. A Shapiro-Wilk test found that not all data are normally distributed. Individual data points and group medians with 95% CI are shown.

**Figure S8.**
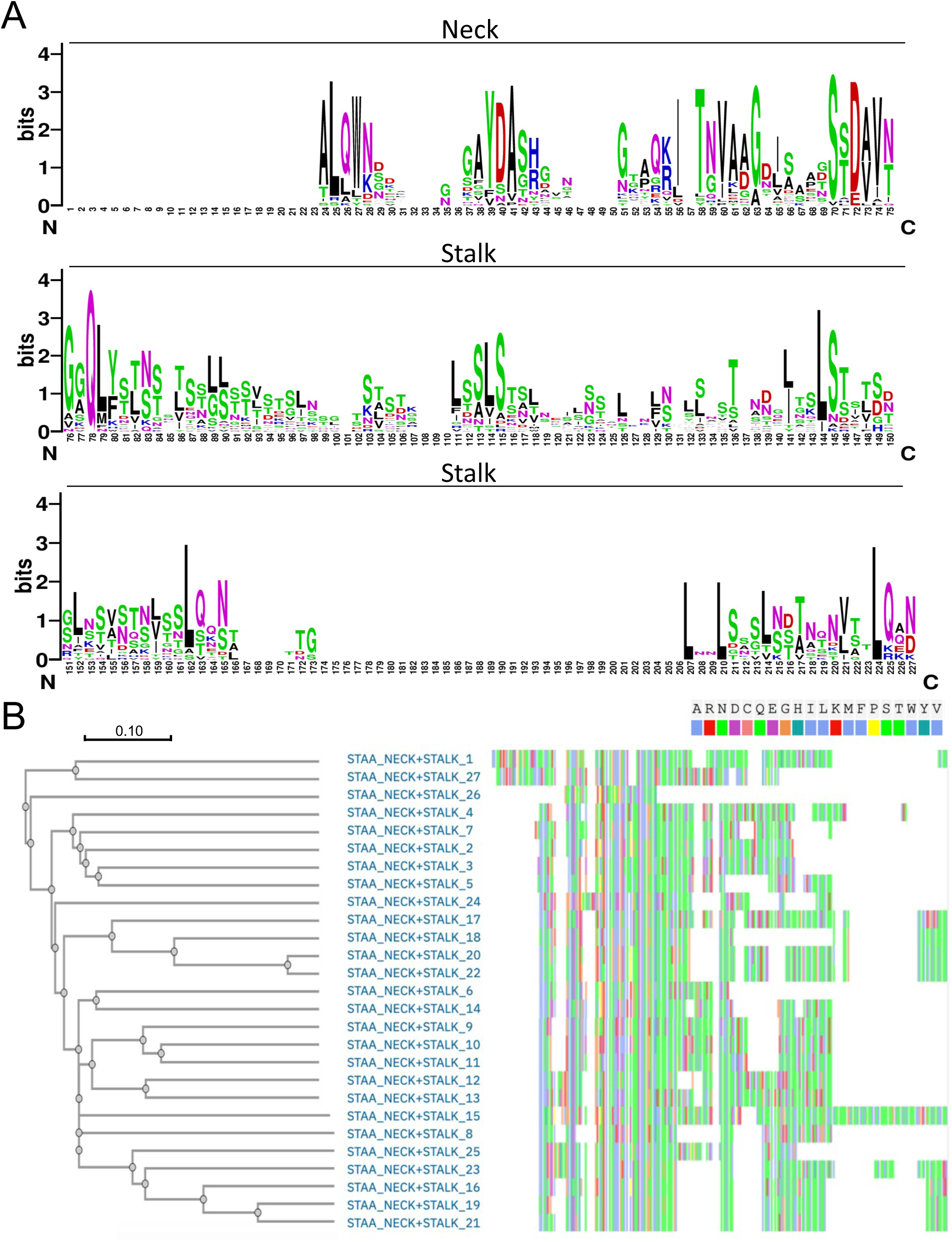
StaA neck + stalk repeats display sequence conservation. A. Sequence logo of residues with high identity among 27 StaA neck (top) and stalk (middle, bottom) repeats, highlighting sequence conservation. X axis: residue position, from N to C terminus. Letter size indicates residue frequency at a given position among all 27 repeats (larger letter = higher frequency of occurrence). Letter color indicates side residue charge [black: hydrophobic; red: negative charge; blue: positive charge; green: polar (no amide); purple: polar (with amide)]. B. Phylogenetic tree of StaA neck + stalk repeats (left), highlighting sequence relatedness. Tree is drawn to scale, as indicated. Residue identity for each repeat is shown (right) to visualize regions of sequence conservation and variability among repeats. Key indicates residue color (right).

**Figure S9.**
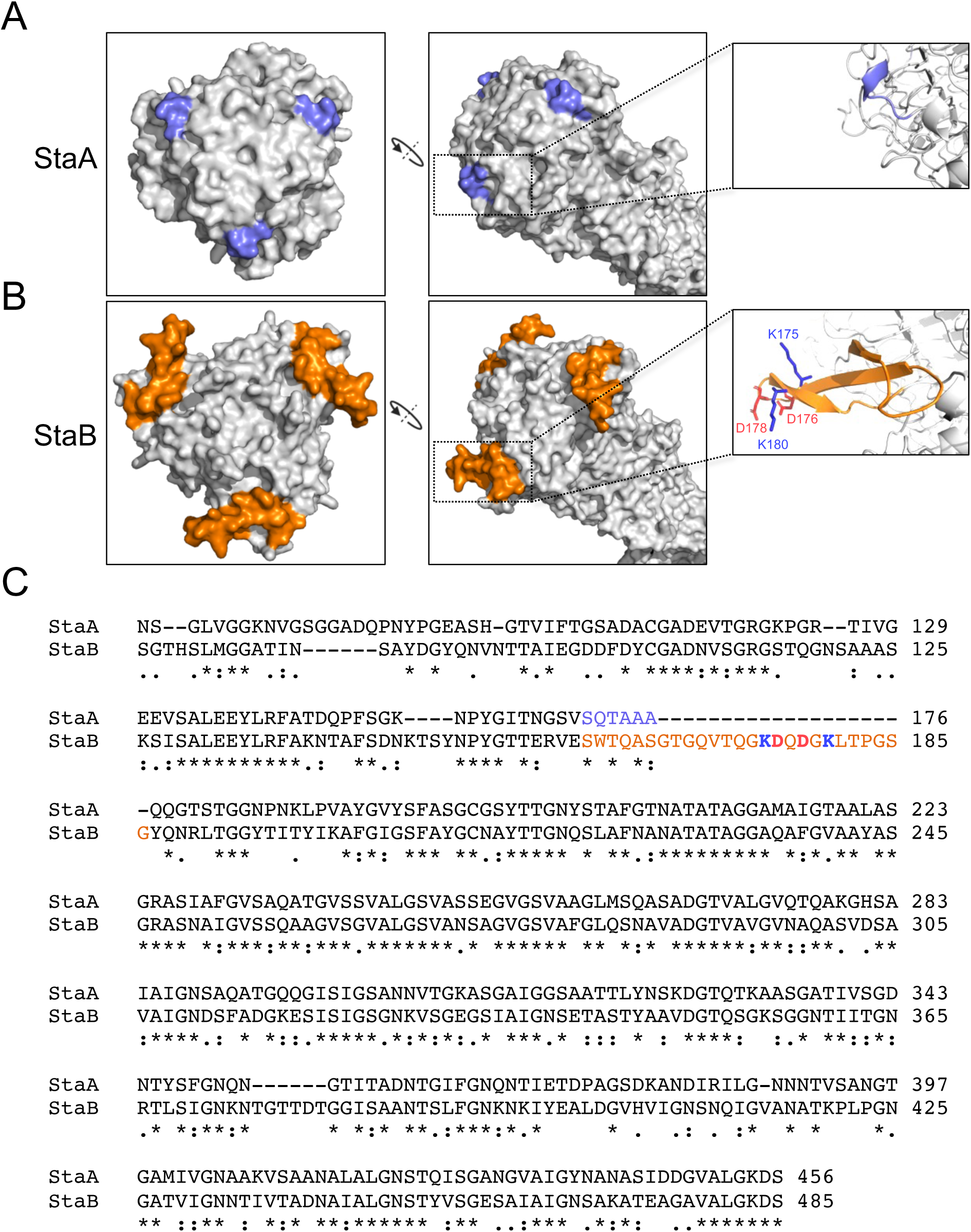
StaB contains a charged head loop absent in StaA. A. Predicted structure of StaA head (left, middle), showing that it lacks a charged head loop (light blue). Inset (right) shows lack of charged loop at the residue level. B. Predicted structure of StaB head (left, middle), showing that it contains a charged head loop sticking into space. Inset (right) shows unsatisfied charged residues in the head loop. Charged residues are labelled by position and colored according to charge (dark blue: positively charged; red: negatively charged). C. Sequence alignment of StaA and StaB head domains, showing regions of conservation and variability. Residue conservation is denoted by symbols below sequences: “*” = Identical residue; “:” = conservative mutation; “.” = semi-conservative mutation. Residue number is indicated at the right side of each row. Gaps are indicated by “-“. Residues present in the head loops shown in panels A and B are indicated by color: Light blue = StaA uncharged head loop; Orange = StaB charged head loop. Residue charge is indicated by color: Dark blue = positively charged; red = negatively charged.

**Figure S10.**
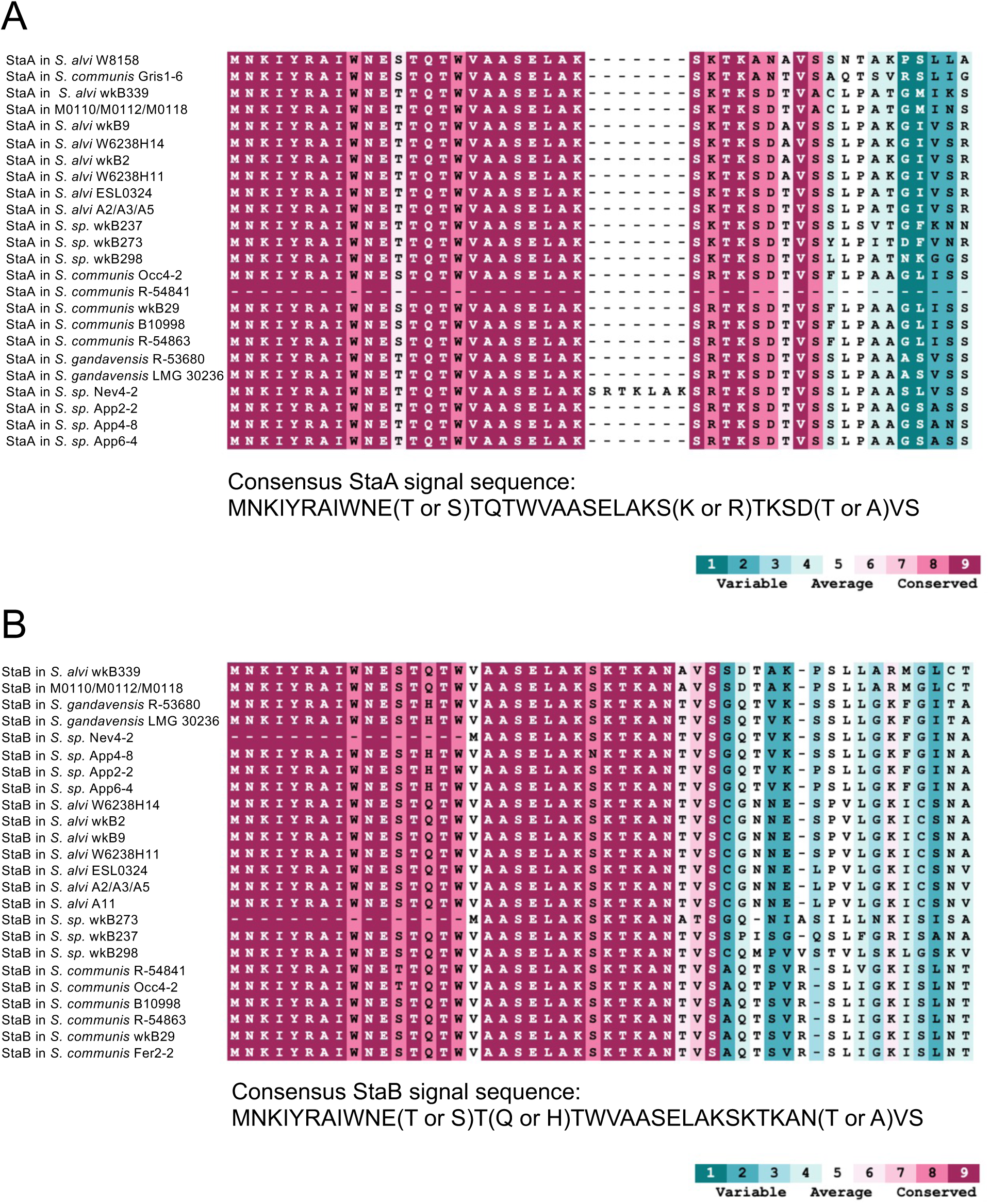
StaA and StaB each have conserved, related signal sequences. A. Alignment and conserved residue analysis of StaA signal sequences across *Snodgrassella* (top), indicating the StaA signal sequence is conserved across the genus. Degree of residue conservation is indicated by color according to key (maroon = more conserved; turquoise = more variable). Gaps are indicated by “-“. The consensus StaA signal sequence is shown (bottom). Variable residues within the consensus sequence are indicated. B. Alignment and conserved residue analysis of StaB signal sequences across *Snodgrassella* (top), indicating the StaB signal sequence is conserved across the genus. Degree of residue conservation is indicated by color according to key (maroon = more conserved; turquoise = more variable). Gaps are indicated by “-“. The consensus StaB signal sequence, shown (bottom), is similar to the consensus StaA signal sequence (panel A). Variable residues within the consensus sequence are indicated.

**Figure S11.**
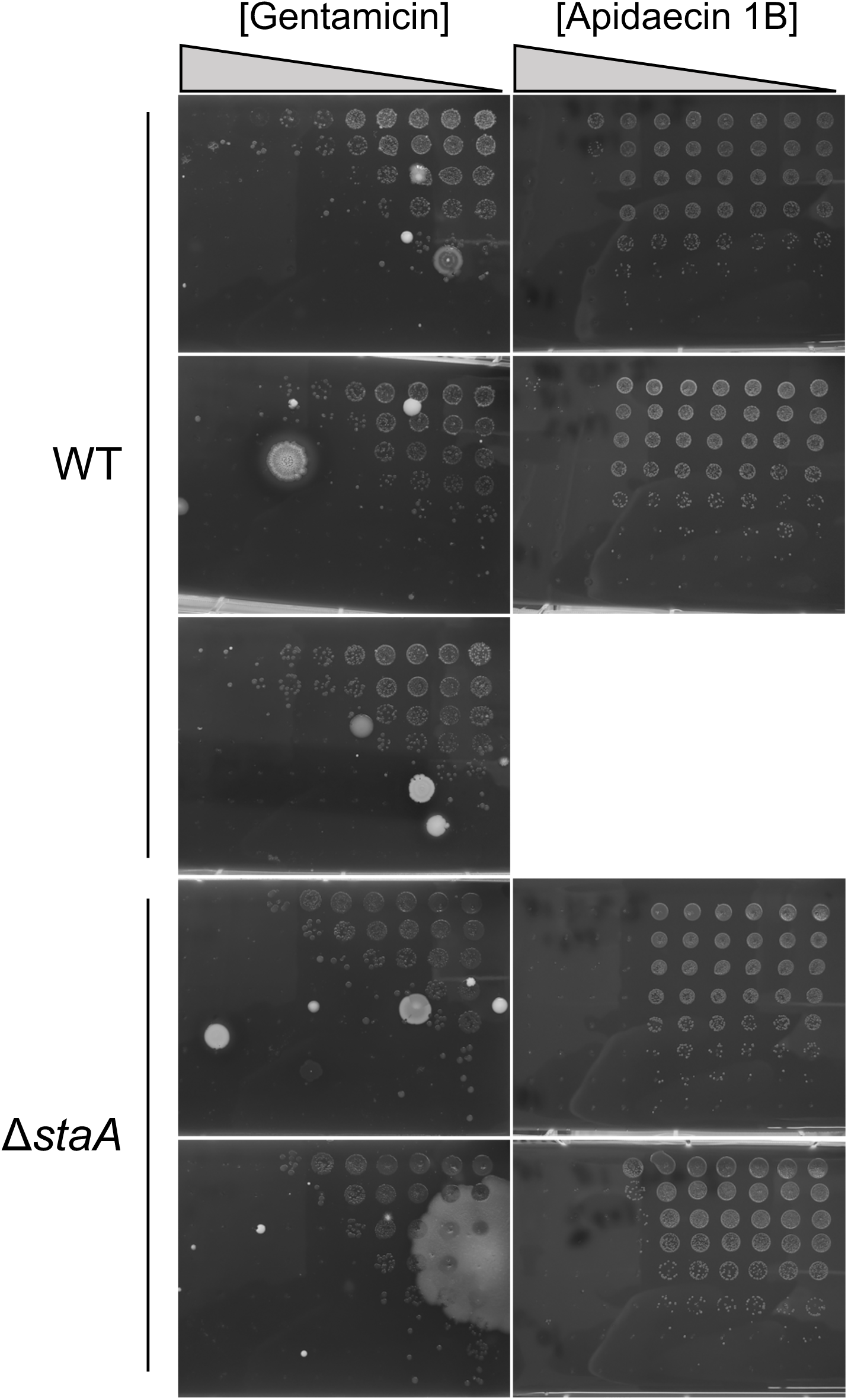
Viability of WT and Δ*staA S. alvi* cells exposed to antibiotic dilution series. Plates of CFU counts of WT and Δ*staA S. alvi* cells grown in 96 well plates and exposed to antibiotics used to generate log_10_ fold change in CFUs/ml plots in Figure 6. For each strain, each row of plates is a biological replicate. For individual plates, each row contains a 2-fold antibiotic dilution series and each column is a 10-fold cell dilution series, to allow for CFU counting.

